# Host Immunity to *Mycobacterium tuberculosis* Infection is Similar in Simian Immunodeficiency Virus (SIV)-infected, Antiretroviral Therapy-treated and SIV-naïve Juvenile Macaques

**DOI:** 10.1101/2022.12.14.520525

**Authors:** Erica C. Larson, Amy L. Ellis, Mark A. Rodgers, Abigail K. Gubernat, Janelle L. Gleim, Ryan V. Moriarty, Alexis J. Balgeman, Yonne K. Menezes, Cassaundra L. Ameel, Daniel J. Fillmore, Skyler M. Pergalske, Jennifer A. Juno, Pauline Maiello, Alexander G. White, H. Jacob Borish, Dale I. Godfrey, Stephen J. Kent, Lishomwa C. Ndhlovu, Shelby L. O’Connor, Charles A. Scanga

**Author notes:** Address correspondence to Erica C. Larson, and Charles A. Scanga.

## Abstract

Pre-existing HIV infection increases tuberculosis (TB) risk in children. Antiretroviral therapy (ART) reduces, but does not abolish, this risk in children with HIV. The immunologic mechanisms involved in TB progression in both HIV-naïve and HIV-infected children have not been explored. Much of our current understanding is based on human studies in adults and adult animal models. In this study, we sought to model childhood HIV/*Mycobacterium tuberculosis* (Mtb) coinfection in the setting of ART and characterize T cells during TB progression. Macaques equivalent to 4-8 year-old children were intravenously infected with SIVmac239M, treated with ART three months later, and coinfected with Mtb three months after initiating ART. SIV-naïve macaques were similarly infected with Mtb alone. TB pathology and total Mtb burden did not differ between SIV-infected, ART-treated and SIV-naïve macaques, although lung Mtb burden was lower in SIV-infected, ART-treated macaques. No major differences in frequencies of CD4+ and CD8+ T cells and unconventional T cell subsets (Vγ9+ γδ T cells, MAIT cells, and NKT cells) in airways were observed between SIV-infected, ART-treated and SIV-naïve macaques over the course of Mtb infection, with the exception of CCR5+ CD4+ and CD8+ T cells which were slightly lower. CD4+ and CD8+ T cell frequencies did not differ in the lung granulomas obtained at necropsy, nor did they differ in the frequency of immune checkpoint and proliferative markers. Thus, ART treatment of juvenile macaques, three months after SIV infection, resulted in similar progression of Mtb and T cell responses compared to Mtb in SIV-naïve macaques.

## Introduction

Pediatric tuberculosis (TB) caused by the bacterium, *Mycobacterium tuberculosis* (Mtb), is a major global health concern. In 2019, around 1.2 million children under the age of 15 fell ill with TB and over 200,000 children died of TB, including children with HIV-associated TB (1). HIV-infected children have higher rates of mortality due to TB than HIV-uninfected children (2). Children account for roughly 10% of HIV-associated TB deaths, which amounted to ∼20,000 lives in 2020 (1, 2). Antiretroviral therapy (ART) reduces TB risk and mortality by suppressing viral replication and restoring CD4+ T cell levels, but TB risk does not completely return to the level seen in HIV-naive children (3–7). Moreover, pediatric TB often manifests differently than adults and disease progression is influenced by age (8, 9). Miliary TB and TB meningitis is more common in infants and young children (< 2 years old), while pulmonary TB is more common in older children (9). HIV infection exacerbates TB disease in children and is associated with greater lung involvement and cavitation regardless of age (10, 11). Given the severity of TB in children, especially those with HIV, there is a clear need to elucidate immune mechanisms underlying TB progression in children as it may help inform diagnostic and treatment strategies.

Much of what is known about pediatric TB is through the lens of human adult studies and adult animal models. However, this overlooks the dynamic nature of the developing, pediatric immune system (12). Throughout childhood, T cell composition is incredibly dynamic and does not stabilize until adulthood (13–16). Rapid accumulation of circulating memory CD4+ and CD8+ T cells occurs during the first few years of life in both children and young nonhuman primates (NHP) (13, 17, 18). Given that CD4+ and CD8+ T cells are critical for Mtb control (19, 20) and the predominant immature nature of CD4+ and CD8+ T cells during the first few years of life, this may be a contributing factor to severe TB disease observed in young children.

However, the role of CD4+ and CD8+ T cells in TB pathogenesis in children is largely understudied. In addition, HIV infection is well-known to cause CD4+ and CD8+ T cell dysfunction through CD4+ T cell depletion and T cell exhaustion (21–24). ART has been shown to restore CD4+ T cell levels and improve CD8+ T cell function, but the immune restoration is incomplete (25–28). Similarly, unconventional T cells, such as γδ T cells, MR1-restricted mucosal-associated invariant T (MAIT) cells, and CD1d-restricted natural killer T (NKT) cells have received little attention in pediatric TB despite their ability to recognize non-peptide Mtb antigens, and may play a possible role in early Mtb control (29–33). Vδ2+ γδ T cells, a subset of γδ T cells which forms T cell receptor heterodimers with Vγ9, and MAITs are virtually absent in early life in humans (16). Moreover, unconventional T cell subsets are depleted during HIV infection and only partially restored by ART (34–36). Whether their role in TB progression differs between HIV, ART-treated children and HIV-naïve children has yet to be thoroughly investigated.

NHP are an excellent model to study TB as they closely recapitulate the immune responses and pathogenesis observed in humans (37, 38). NHP are also susceptible to SIV, a close relative of HIV, which results in HIV-like disease progression and AIDS development in some macaque species (39). Previously, in adult Mauritian cynomolgus macaques, we found that Mtb coinfection of ART-naïve, SIV-infected animals had worsened TB disease compared to SIV-naïve macaques (40), in alignment with studies in humans (41–43). In a separate study, we observed granulomas obtained from SIV/Mtb coinfected macaques early in the course of Mtb infection had immunologic differences compared to animals infected with Mtb alone, such as fewer CD4+ T cells, more CD8+ T cells, and elevated frequencies of PD-1+ and TIGIT+ T cells, indicative of chronic immune activation (44). Although these studies inform our understanding of TB immunity in coinfected adults, very few NHP studies to date have modeled pediatric TB (45–47).

This is the first study to characterize CD4+ and CD8+ T cell populations over the course of infection using an NHP model of pediatric TB and HIV/Mtb coinfection. Juvenile macaques, ∼1-2 years of age (equivalent to 4-8 years in humans), were either infected with Mtb alone or were infected with SIV, treated with ART, and then coinfected with Mtb. We found very few differences in TB disease progression and Mtb burden between SIV-infected, ART-treated and SIV-naïve macaques. While we did observe immunological changes following SIV infection, such as fewer CD4+ T cells and more CD8+ T cells in airways, these returned to pre-SIV levels following ART initiation. The frequencies of CD4+ and CD8+ T cells in airways remained similar between SIV-infected, ART-treated and SIV-naïve macaques 8 weeks after Mtb infection, although. Frequencies of unconventional T cell subsets (Vγ9+ γδ T cells, MAIT cells, and NKT cells) did not differ over the course of SIV nor between SIV-infected, ART-treated and SIV-naïve macaques following Mtb infection. Similarly, T cell composition of granulomas did not differ between the two groups. Thus, juvenile macaques treated with ART within 3 months of SIV infection appear to experience similar TB progression and mount similar T cell responses to Mtb coinfection as animals infected with Mtb alone.

## Results

### T cell subsets and phenotype of CD4+and CD8+ T cells in airways do not differ, except for CCR5, between SIV-infected, ART-treated and SIV-naïve juvenile macaques

Ten juvenile macaques were infected intravenously with SIVmac239M and, as expected, peak viremia was detected approximately 10 days post infection followed by viral load reduction and establishment of set-point viremia, which varied widely among animals. Three months after infection, ART was initiated in all animals and reduced viremia to below the limit of detection (Figure 1). Plasma viral load remained undetectable in all animals after ART initiation, with the exceptions of 34519 and 34619 which had transient viremia that did not exceed 10^3^ viral copies (Figure 1). All 10 SIV-infected, ART-treated animals, as well as 10 age-matched SIV-naïve animals, were then infected with approximately 5-11 colony forming units (CFU) of a barcoded Mtb (Table S1). ART continued to suppress viral replication after Mtb challenge in SIV-infected animals (Figure 1).

**Figure 1.**
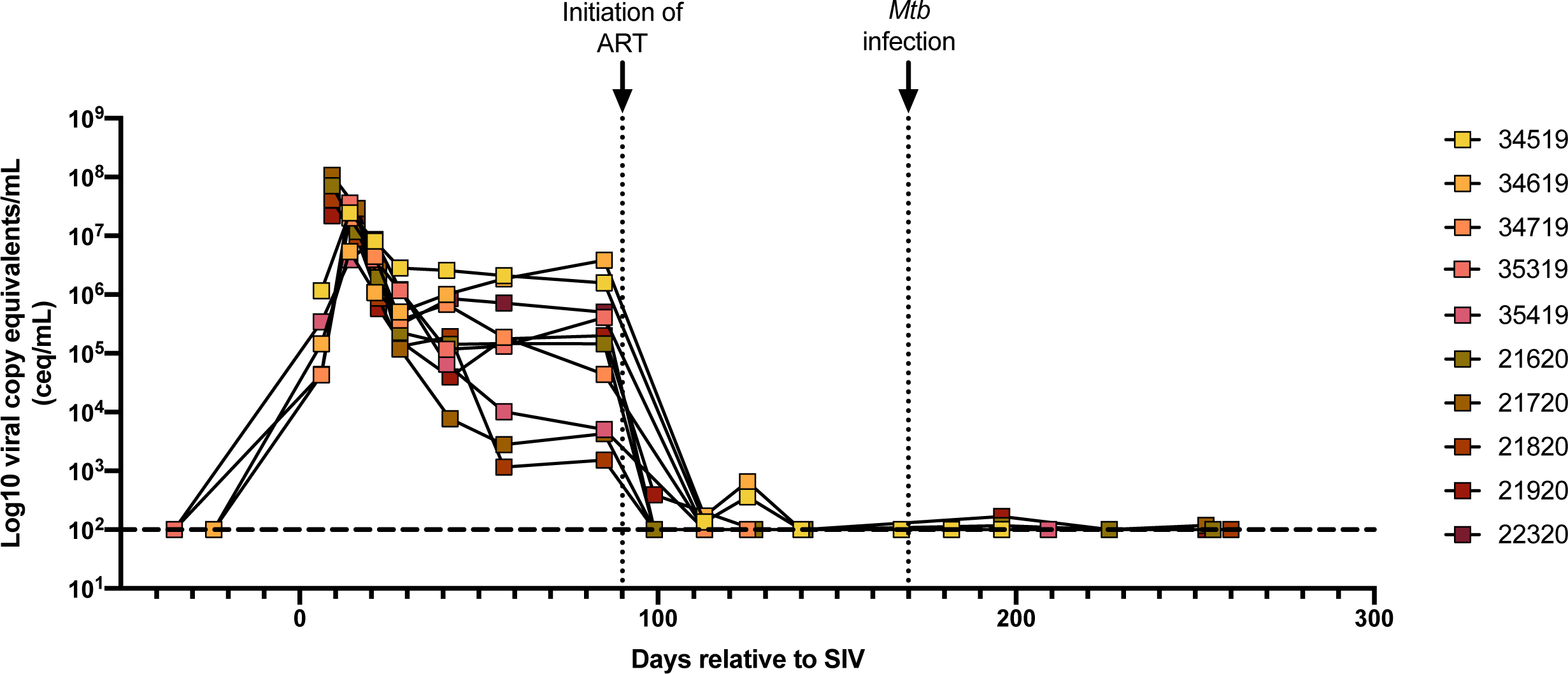
Plasma viremia of SIV-infected, ART-treated macaques over the course of SIV and Mtb coinfection. Plasma viral copy equivalents were determined by qRT-PCR. Each point indicates an individual animal. Horizontal dashed line represents the limit of detection.

Mtb is predominantly transmitted through inhaled droplets and first encounters host immune cells when deposited in the airways. Thus, we assessed T cell composition and characterized T cell phenotype in the airways of the animals during SIV infection and ART treatment by flow cytometric analysis of cells recovered by bronchoalveolar lavage (BAL). The composition of T cell subsets dramatically shifted following SIV infection, with a notable decrease in the CD4+ proportion (blue) and a corresponding increase in the CD8+ proportion (red) (top row, Figure 2A). The T cell composition shifted again following ART, returning to proportions similar to those observed prior to SIV infection (Figure 2A). T cell proportions were similar between SIV-infected, ART-treated macaques (top row) and macaques infected with Mtb alone (bottom row), both prior to and 8 weeks after Mtb infection (Figure 2A). These changes in T cell subsets also were noted when data are presented as frequencies of the total CD3+ population (Figure 2B-J). We observed a significant decline in CD4+ T cells with a concomitant rise in CD8+ T cell frequencies in the airways following SIV infection and a return to pre-SIV frequencies after initiating ART (Figure 2B, C). At the time of Mtb coinfection, CD4+ T cell frequencies in BAL were similar between SIV-infected, ART-treated animals and SIV-naïve controls and did not change appreciably after Mtb infection (Figure 2B). CD8+ T cells exhibited a subtle, but significant, decline in both groups following Mtb coinfection (Figure 2C). CD4+CD8+ T cell frequencies increased following Mtb infection in both groups (Figure 2D), which has been reported previously (48). We did not observe significant differences in CD3+CD4-CD8-T cells or consistent changes in unconventional T cell subsets, including γδ T cells (Vγ9+), NKT cells (CD1d tetramer+), MAIT cells (MR1 tetramer+ Vα7.2+), and MAIT-like cells (MR1 tetramer+Vα7.2-) between the two groups following Mtb coinfection (Figure 2E-I).

**Figure 2.**
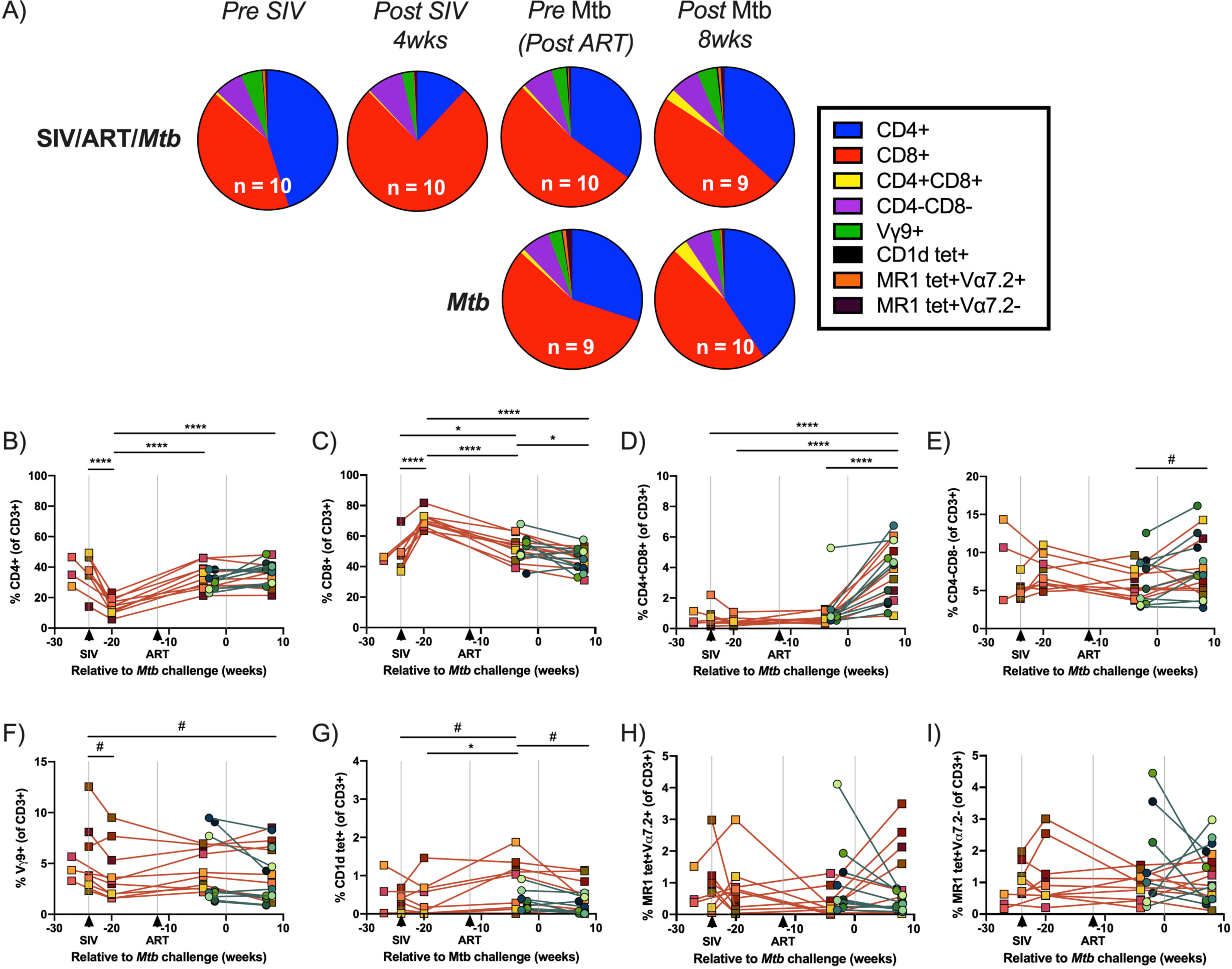
Composition of T cell subsets in airways over time. BALs were collected over the course of SIV and Mtb infection. A) Proportions of T cell subsets by median event counts. B-J) Frequencies of T cell subsets over time relative to CD3+ gate. Individual animals indicated by symbols. SIV-infected, ART-treated Mtb coinfected macaques are indicated by orange lines and macaques infected with Mtb alone are indicated by teal lines. A mixed statistical model was performed to determine significance with fixed effects (time and group) and random effects (individual animals). Bars with asterisks indicate significance between time points; # 0.05 < p < 0.1, * p < 0.05, and **** p < 0.0001.

We did observe some phenotypic changes in CD4+ and CD8+ T cells in airways over the course of the study. There was a transient spike in proliferation, as measured by ki-67, in CD4+ and CD8+ T cells at 4 weeks post-SIV infection which then subsided following ART to levels similar to those observed in uninfected macaques (Figure S1A, B). CD4+ and CD8+ T cell proliferation was previously shown to correspond to the rise in viral replication during acute SIV infection (49–51). Following Mtb infection, the frequency of PD-1+ CD4+ T cells, but not PD-1+ CD8+ T cells, declined significantly (Figure S1C, D). The frequency of TIGIT+ CD4+ and CD8+ T cells fluctuated, with notable increases in frequency of TIGIT+ CD4+ T cells 4 weeks post SIV infection and of TIGIT+ CD8+ T cells following Mtb infection (Figure S1E, F). A small, but significant, drop in the frequency of CXCR3+ CD8+ T cells, but not CD4+ T cells, was noted following Mtb coinfection (Figure S1G, H). CCR6-expressing CD4+ and CD8+ T cells significantly increased in frequency in both groups following Mtb coinfection (Figure S1I, J). As CCR6 mediates cell migration during inflammation and immune response (52), this increased frequency of CCR6+ CD4+ and CD8+ T cells most likely indicates enhanced trafficking to the airways in response to Mtb (53).

The frequency of CCR5, on the other hand, significantly declined in CD8+ T cells, and to a lesser extent in CD4+ T cells (p = 0.0534), in SIV-infected, ART-treated macaques following Mtb coinfection (Figure 3A, B). CCR5 is a coreceptor utilized by HIV/SIV for infection of CD4+ T cells and is closely linked to T cell loss (54, 55). This decline was also reflected by a loss of absolute CCR5+ CD4+ and CD8+ T cell numbers in the airways (Figure 3C). SIV-infected, ART-treated animals had significantly fewer CD4+ and CD8+ T cells in their airways (∼0.39 and 0.40 log decrease, respectively) than compared SIV naïve animals (Figure 3C). Given the role of CD4+ and CD8+ T cells in Mtb control, we were interested in whether this subtle loss of conventional T cells in the airways impacted overall TB progression.

**Figure 3.**
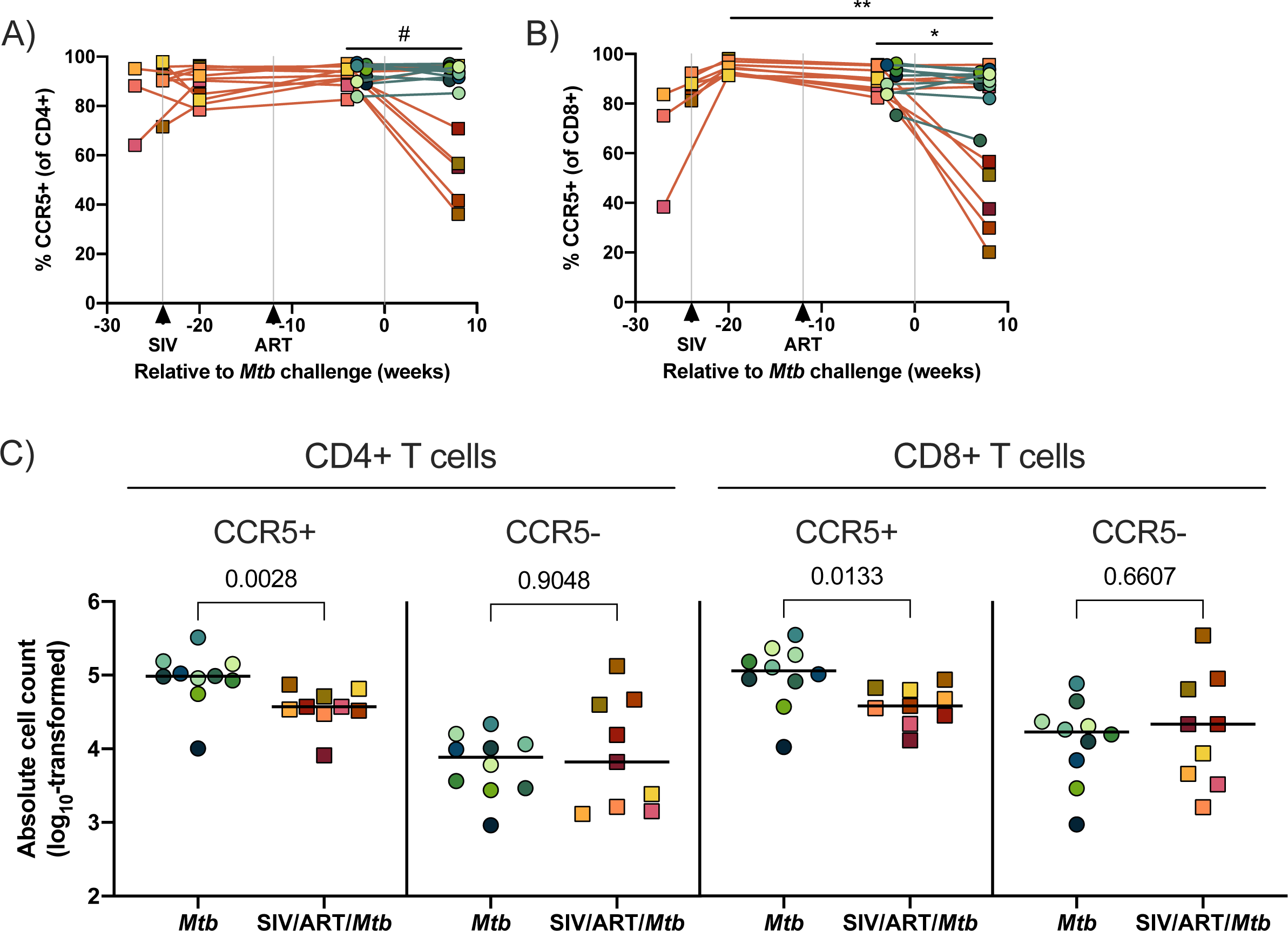
CCR5+ CD4+ and CD8+ T cells in airways at 8 weeks post Mtb infection. BAL cells were collected over the course of SIV and at 8 weeks post Mtb infection and stained for flow cytometry. A&B) Individual animals indicated by symbols. SIV-infected, ART-treated Mtb coinfected macaques are indicated by orange lines and macaques infected with Mtb alone are indicated by teal lines. A mixed statistical model was performed to determine significance with fixed effects (time and group) and random effects (individual animals). Bars with asterisks indicate significance between time points; # 0.05 < p < 0.1, * p < 0.05, and ** p < 0.01. A) Frequencies of CCR5+ CD4+ T cells over the course of the study. B) Frequencies of CCR5+ CD8+ T cells over the course of the study. C) Absolute CCR5+/-CD4+ and CD8+ T cells in airways at 8 weeks post Mtb. Absolute cell counts were calculated from the hemacytometer cell count. Individual samples indicate individual animals and bars indicate group medians. Mann Whitney U tests was performed to determine significance between groups. P-values are shown.

### No difference in lung inflammation or Mtb dissemination

We previously reported a dramatic increase in lung inflammation and Mtb dissemination in SIV-infected, ART-naïve adult macaques between 4-8 weeks post Mtb compared to adult macaques infected with Mtb alone, indicating a loss of Mtb control in the SIV-infected group (40). Here, we used PET/CT to determine whether lung inflammation and Mtb dissemination differed over the course of Mtb infection in our juvenile SIV-infected, ART-treated macaques compared to juvenile macaques infected with Mtb alone. We measured FDG uptake, a surrogate for inflammation, to quantify lung inflammation and to enumerate granulomas over time as a measure of Mtb dissemination (Figure 4). SIV-infected, ART-treated macaques coinfected with Mtb did not differ from macaques infected with Mtb alone in terms of total lung FDG activity over the course of Mtb infection (Figure 4A). Similarly, both groups exhibited similar numbers of granulomas over the Mtb infection course (Figure 4B). One SIV-infected, ART-treated animal had rapidly progressive TB disease and reached humane endpoint just 6 weeks post-Mtb coinfection and is represented as a single datapoint at 4 weeks. When measured at the final time point, there were no significant differences in lung inflammation (p = 0.2415) or the number of granulomas (p = 0.4601) between SIV-infected, ART-treated macaques coinfected with Mtb and macaques infected with Mtb alone (Figure 4C, D), indicating similar TB progression in the two groups.

**Figure 4.**
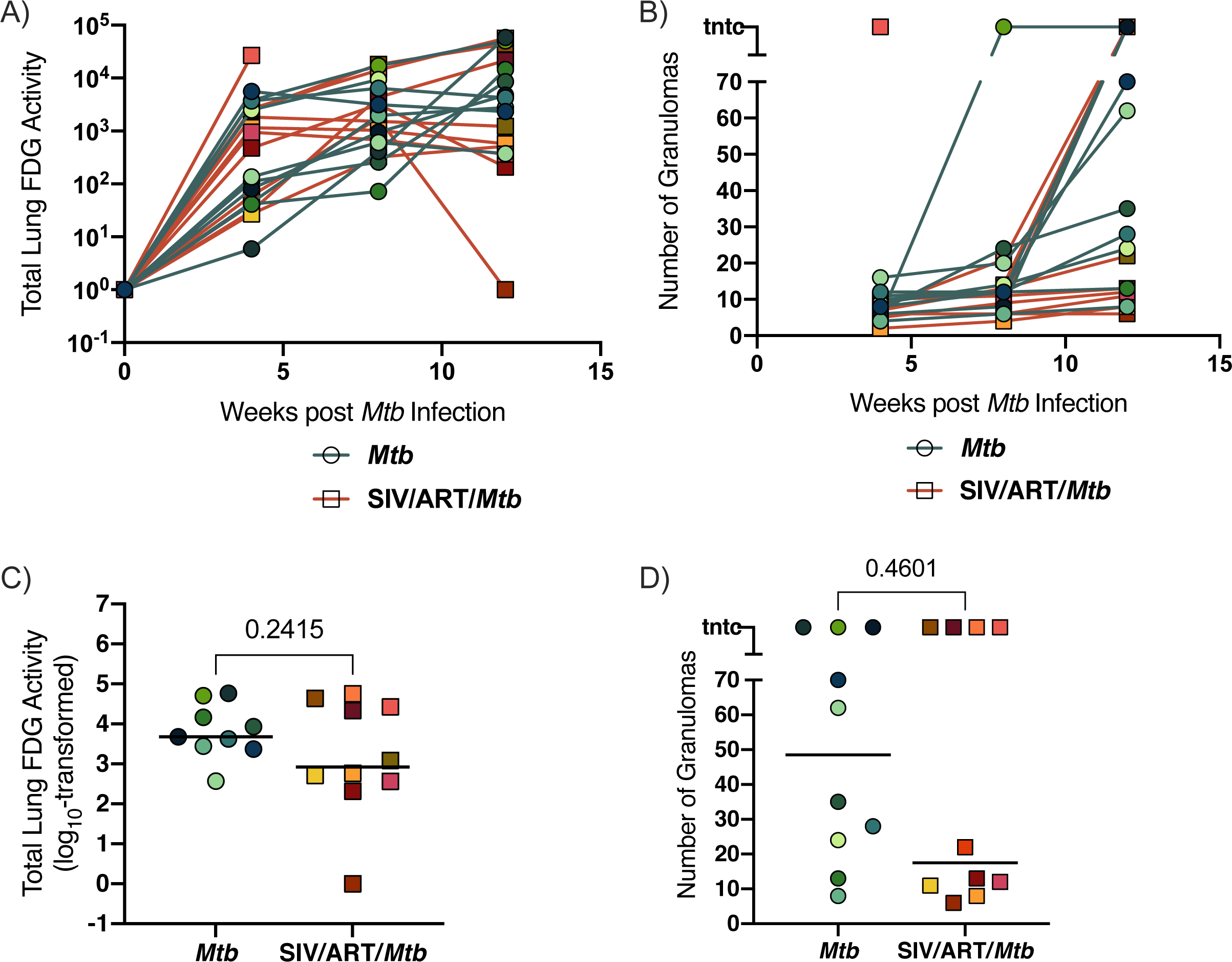
TB progression by PET/CT imaging. A) Lung FDG activity over time. SIV-infected, ART-treated Mtb coinfected macaques (orange lines) and macaques infected Mtb alone (teal lines). Individual animals are shown. B) The number of granulomas identified by PET-CT at 4, 8, and 12 weeks (pre-necropsy). SIV-infected, ART-treated Mtb coinfected macaques (orange lines) and macaques infected Mtb alone (teal lines). Individual animals are shown. C) Total lung FDG activity at time of necropsy. Bars indicate medians of group. An unpaired t test of group medians was performed. *P*-value is shown. D) The number of granulomas identified at necropsy (12 wks *p.i. Mtb*). Bars indicate medians of group. A Mann Whitney U test of group medians was performed. *P*-value is shown.

### TB pathology, bacterial burden, and bacterial dissemination were similar in SIV-infected, ART-treated and SIV-naïve juvenile macaques, except for lung CFU

Following Mtb infection, erythrocyte sedimentation rates were normal in both SIV-infected, ART-treated macaques and SIV-naïve macaques while culturable bacilli were variably detected in BAL and gastric aspirates from both groups (Table S1). At necropsy, we used an established scoring system to assess total TB pathology across several tissue compartments: lungs, thoracic lymph nodes, and extrapulmonary sites (56). While the individual pathology scores varied widely, there were no significant differences in the group medians between SIV-infected, ART-treated macaques coinfected with Mtb and macaques infected with Mtb alone (Figure 5A-D).

**Figure 5.**
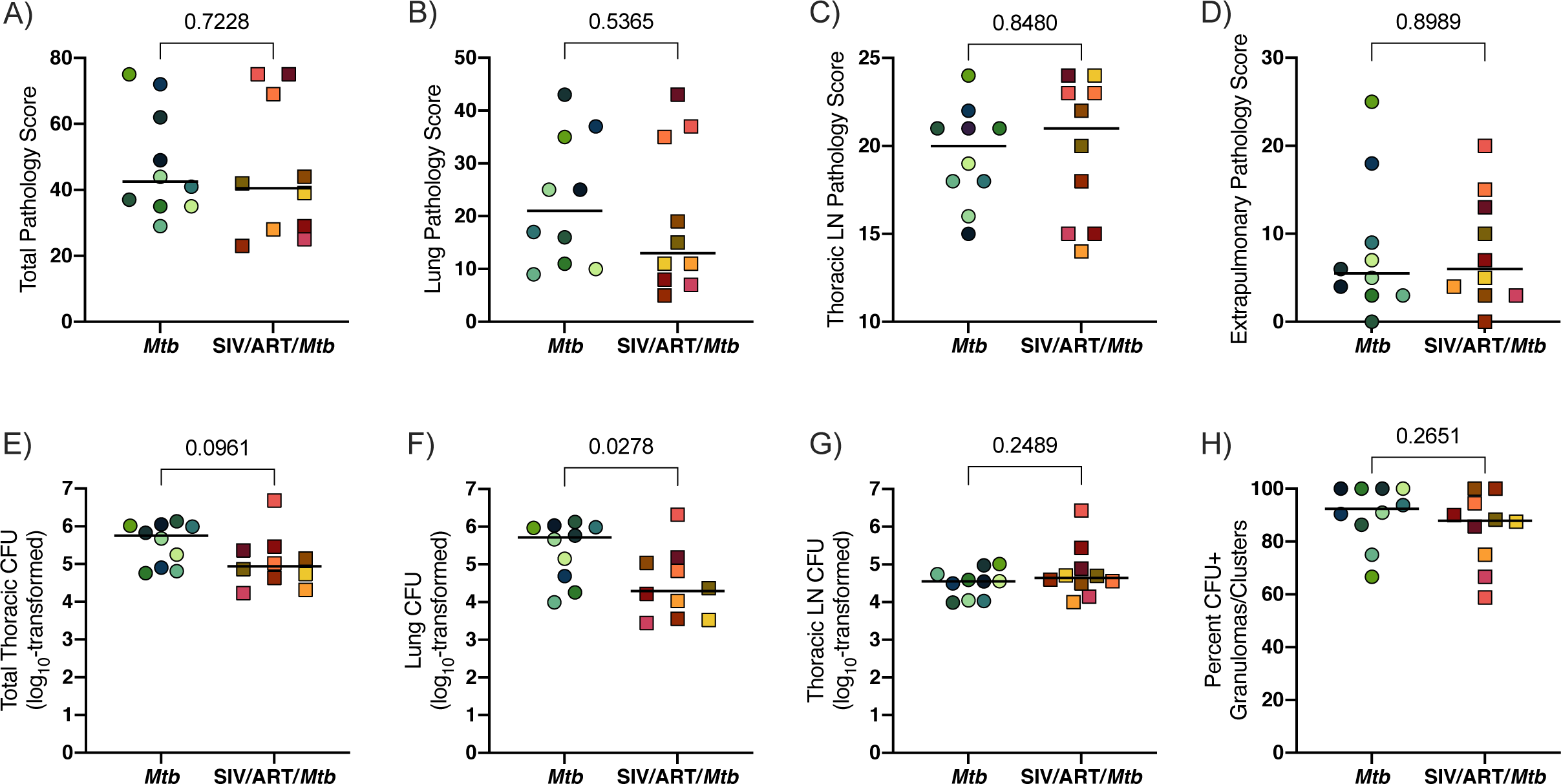
TB pathology and CFU at 12 weeks post Mtb infection. Bars indicate medians of group. Unpaired t tests of group medians were performed. *P*-values are shown. A) Total pathology score. B) Lung pathology score. C) Thoracic lymph node pathology score. D) Extrapulmonary pathology score. E) Total thoracic CFU (lung CFU + thoracic lymph nodes CFU). F) Lung CFU. G) Thoracic lymph nodes CFU. H) Percent CFU+ granulomas or clusters.

We plated tissue samples for CFU to determine Mtb burden. Somewhat surprisingly, the median total thoracic burden of Mtb, comprised of CFU from both lung and thoracic lymph nodes, was slightly lower in SIV-infected, ART-treated macaques compared to macaques infected with Mtb alone although this difference did not reach statistical significance (p = 0.0961; Figure 5E). Interesting, the bacterial load was significantly lower when only the lungs were considered (p = 0.0278; Figure 5F). In contrast, the median bacterial load in thoracic lymph nodes between the groups was similar (p = 0.2489; Figure 5G). Culture-negative lung granulomas identified at necropsy were considered to have been sterilized by the host. Both groups had similar percentages of lung granulomas with culturable bacilli, indicating comparable capacity to eliminate viable Mtb in SIV-infected, ART-treated animals and those that were infected with Mtb alone (Figure 5H).

To assess Mtb dissemination, Mtb DNA was isolated from CFU+ tissue samples and the number and distribution of uniquely tagged bacilli was quantified across individual animals and tissue types. We did not find differences in the median number of uniquely tagged bacilli per animal between cohorts (Figure S2A). The number of uniquely tagged bacilli identified in granulomas and thoracic lymph nodes did not differ between SIV-naïve and SIV-infected, ART-treated, animals (Figure S2B). However, consistent with previous reports of lymph node seeding from multiple granulomas (57, 58), we observed a significantly higher number of uniquely tagged bacilli in thoracic lymph nodes when compared to granulomas. This was the case for both SIV-naïve (p = 0.0397) and SIV-infected, ART-treated animals (p = 0.0002) indicating similar dissemination between these groups.

We also analyzed the bacterial load in individual granulomas from each animal. The number of culturable Mtb within individual granulomas varied widely both within individual animals as well as between animals from each group (Figure 6A). However, there was no significant difference in the median CFU of individual granulomas between those from SIV-infected, ART-treated macaques coinfected with Mtb and those from macaques infected with Mtb alone (p = 0.2047; Figure 6B). Thus, both the severity of TB, Mtb bacterial load, and Mtb dissemination were similar in SIV-naïve and SIV-infected, ART-treated animals.

**Figure 6.**
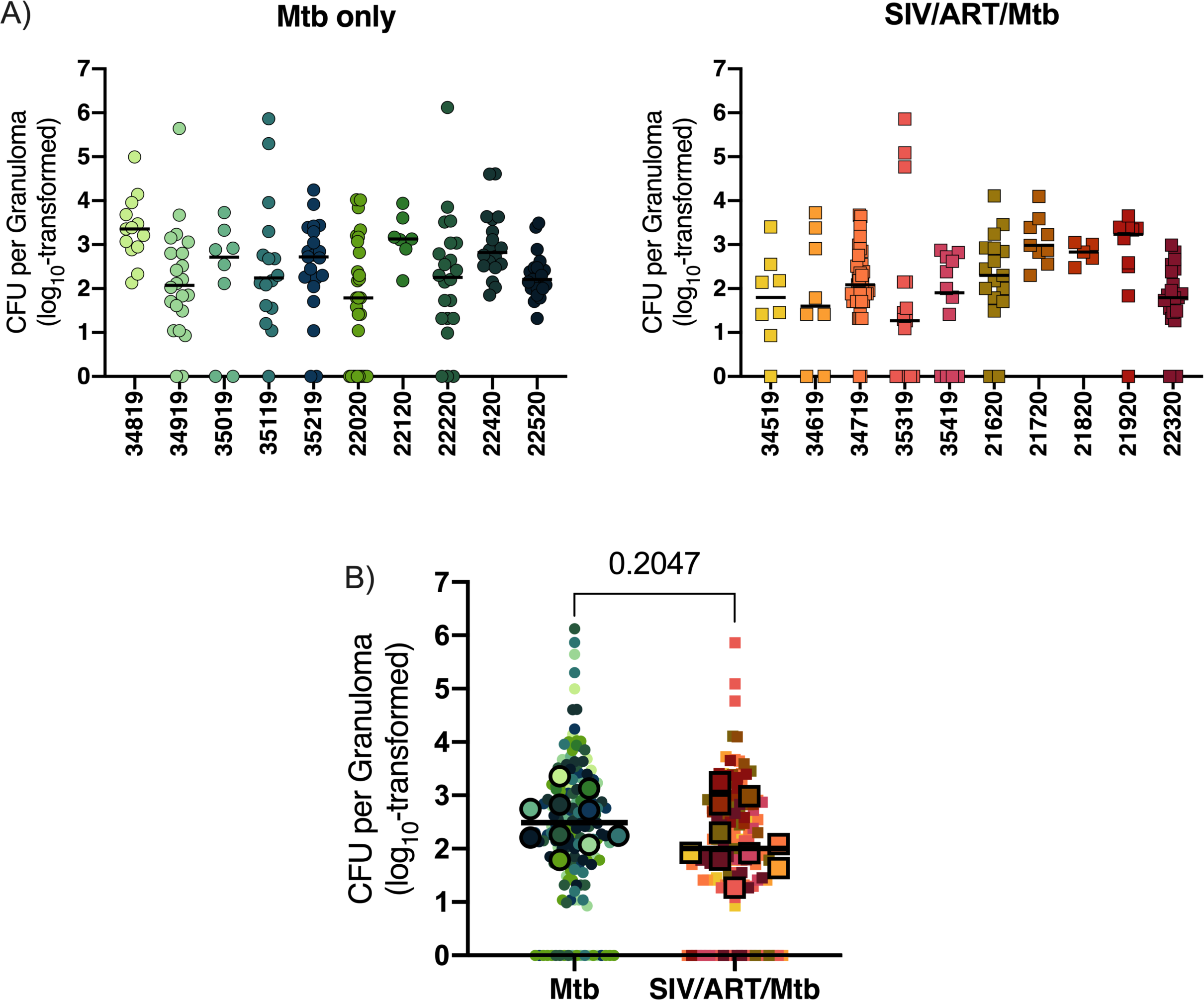
Bacterial burden in individual granulomas. A) CFU per granuloma by animal. Symbols represent CFU from individual granulomas. Bars indicate median CFU per animal. Mtb only animals (left panel) and SIV/ART/Mtb animals (right panel). B) Combined CFU for Mtb only and SIV/ART/Mtb groups. Outlined symbols indicate median per animal and non-outlined symbols indicate individual granulomas. Bars indicate median CFU per animal per group and an unpaired t test was performed to determine significance. The p-value is shown.

### Cytokine responses to Mtb-specific antigens were similar in the lungs of SIV-infected, ART-treated macaques coinfected with Mtb and macaques infected with Mtb alone

Cell suspensions were prepared from lung tissue without apparent granulomas irrespective of whether the tissue would be determined to have culturable Mtb. The cells were stimulated with Mtb whole cell lysate to assess CD4+ and CD8+ T cell responses to Mtb-specific antigens in the lung (Figure 7). There were no differences in IFNγ or TNF production between SIV-infected, ART-treated macaques coinfected with Mtb and macaques infected with Mtb alone in either CD4+ and CD8+ T cells (Figure 7A-D), indicating similar Mtb-specific cytokine responses between the two groups.

**Figure 7.**
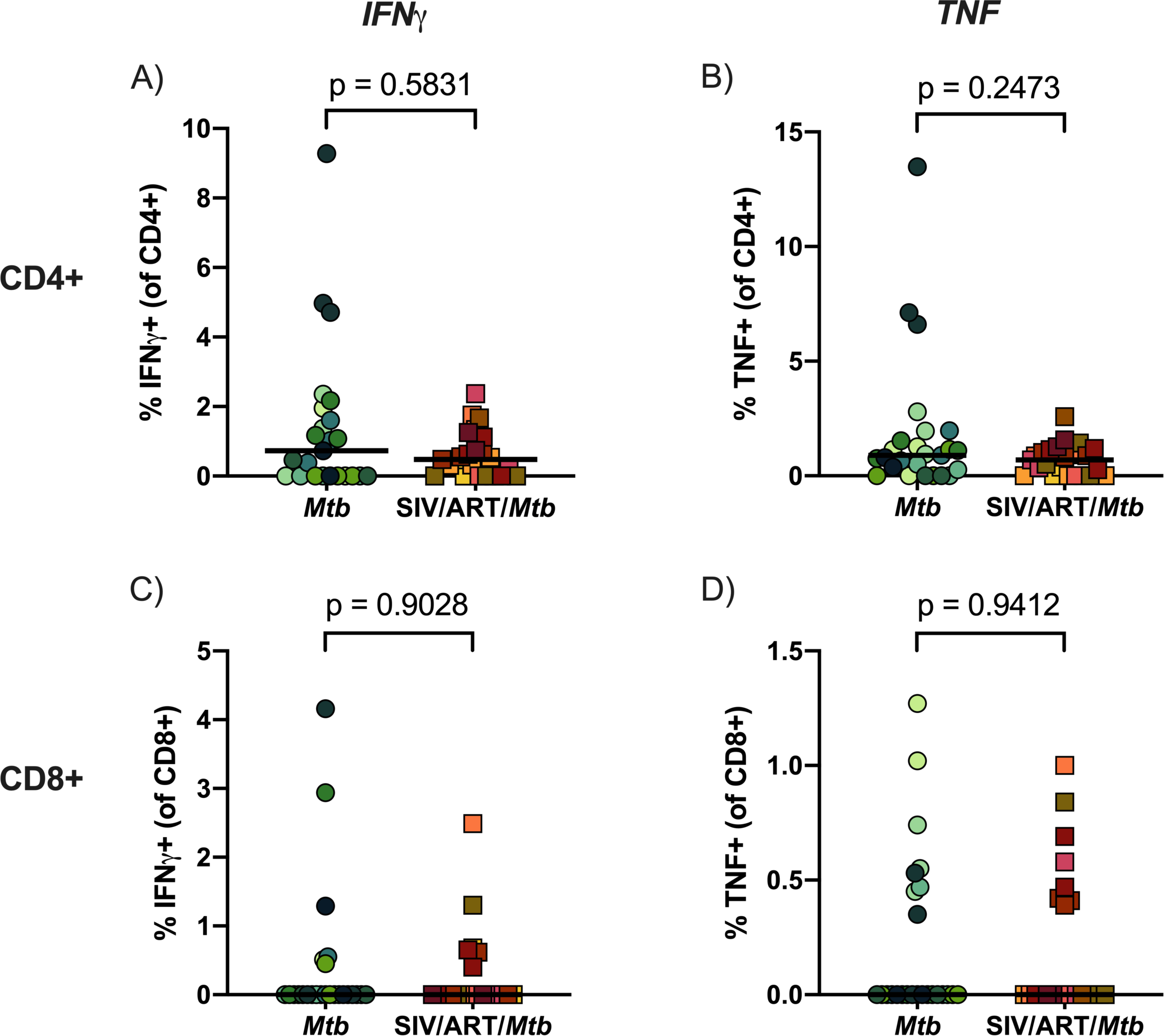
Cytokine responses in lung tissue. Lung tissue was stimulated with H37Rv whole cell lysate for 14 hours. Cytokine production was corrected for against unstimulated controls. Bars indicate medians of pooled data. Mann Whitney U tests were performed to determine significance. *P*-values are shown. A) IFNγ production in CD4+ T cells. B) TNF production in CD4+ T cells. C) IFNγ production in CD8+ T cells. D) TNF production in CD8+ T cells.

### Granuloma composition and phenotype did not differ between Mtb-infected SIV-infected, ART-treated and SIV-naïve macaques

Granulomas are the hallmark of TB disease. While normal granuloma formation is known to be affected in HIV/Mtb coinfected adults (59, 60), very little is known about TB granuloma formation dynamics and composition in children coinfected with HIV, regardless of whether they are treated with ART or not. Histopathology of excised granulomas was assessed by an experienced veterinary pathologist and no generalizable histological differences could be determined in the granulomas from SIV-naïve and SIV-infected, ART-treated animals (data not shown).

We used flow cytometry to compare the cellular composition of TB granulomas in lungs of SIV-infected, ART-treated macaques with those in SIV-naïve macaques. We did not observe any difference in overall T cell composition (Figure 8A) or in frequencies of T cell subsets (Figure 8B-H), nor did T cell frequencies differ in other tissue compartments, including lymph nodes, spleen, and blood (Figure S3).

**Figure 8.**
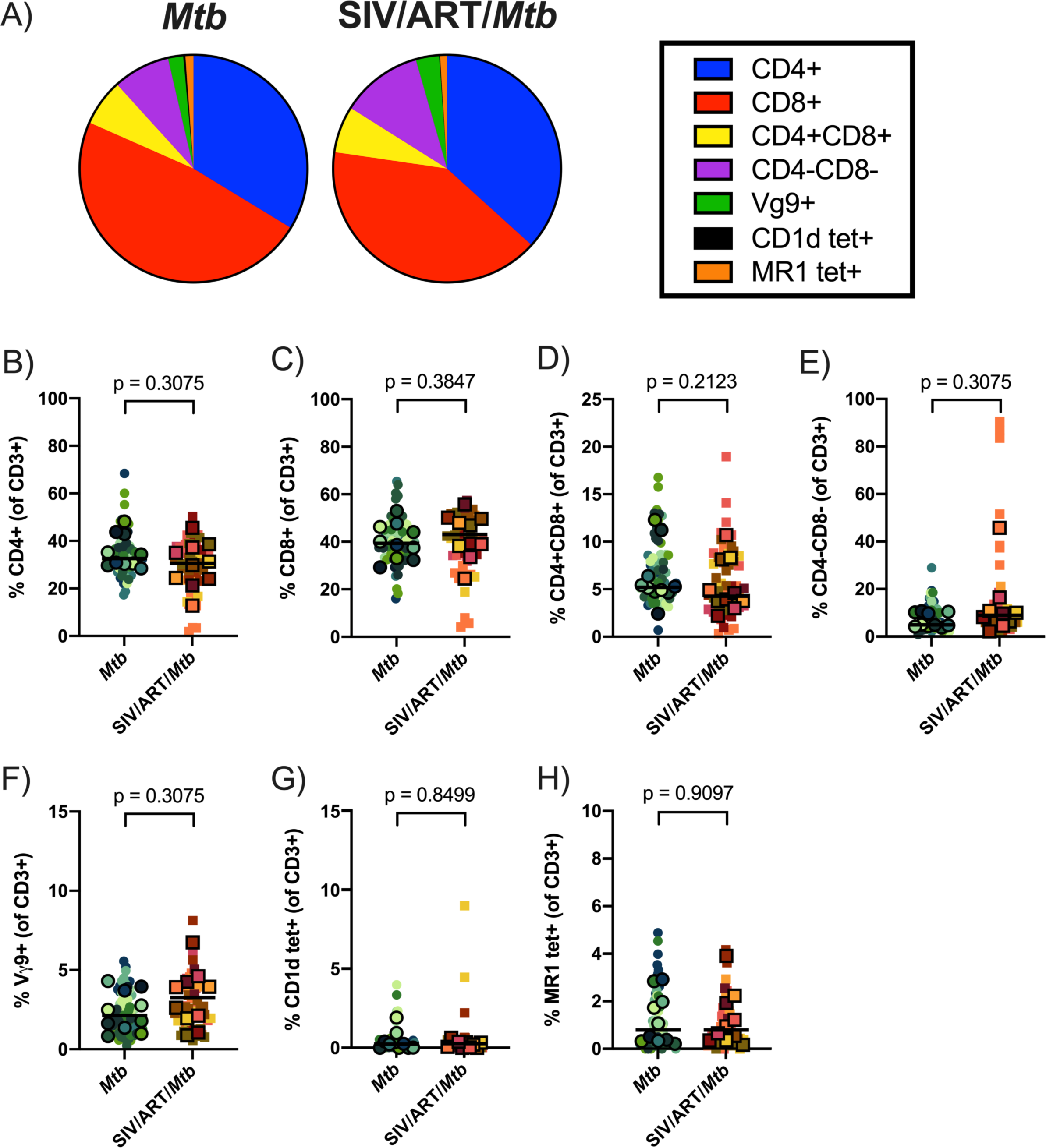
T cell subsets in granulomas. A) Proportion of T cell subsets in individual granulomas by median event counts. B-H) Frequencies of T cell subsets in granulomas relative to CD3+ gate. B) CD4+ T cells. C) CD8+ T cells. D) CD4+CD8+ T cells. E) CD4-CD8-T cells. F) Vγ9+ T cells. G) CD1d tet+ T cells. H) MR1 tet+ T cells. Outlined symbols indicate median per animal and unlined symbols indicate individual samples. Bars indicate group medians. Wilcoxon tests of group medians were performed to determine significance. *P*-values are shown.

Lastly, we investigated the frequency of CCR5 and several other phenotypic markers (PD-1, TIGIT, and ki-67) on CD4+ and CD8+ T cells isolated from lung granulomas (Figure 9 and 10). Unlike the airways, CCR5+ CD4+ and CD8+ T cells did not significantly differ between SIV-infected, ART-treated macaques and macaques infected with Mtb alone (Figure 9A, B). For the other phenotypic markers (Figure 10), there were no significant differences between the two groups apart from TIGIT+ CD4+ T cells (p = 0.0647; Figure 10B), which trended slightly higher in frequency in SIV/ART/Mtb macaques, and ki-67+ CD8+ T cells, which were significantly more frequent in granulomas from SIV-naïve animals (p = 0.0092; Figure 10F). However, the difference in the median frequency of ki-67+ cells between the two groups was incredibly low (< 0.12% of CD8+ T cells in both groups) and warrants caution about ascribing biological significance to this result. Together, these data demonstrate that CD4+ and CD8+ T cells in granulomas from SIV-infected, ART-treated macaques are quite similar to those from macaques infected with Mtb alone.

**Figure 9.**
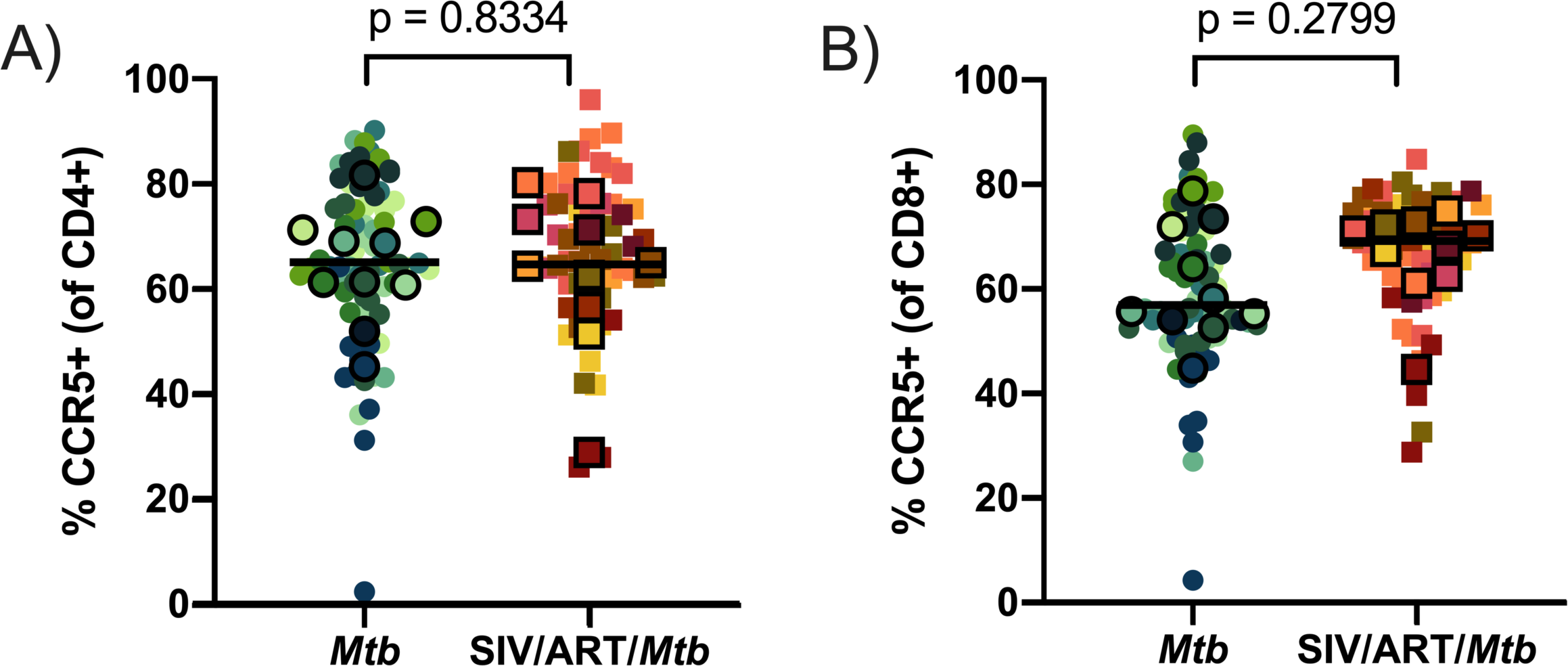
CCR5 frequency of CD4+ and CD8+ T cells isolated from granulomas. Frequencies of CCR5+ CD4+ and CD8+ T cells in granulomas. Outlined symbols indicate median per animal and unlined symbols indicate individual samples. Bars indicate group medians. A) CCR5+ CD4+ T cells. An unpaired t test was performed of group medians to determine significance. B) CCR5+ CD8+ T cells. A Mann Whitney U test of group medians was performed to determine significance. *P*-values are shown.

**Figure 10.**
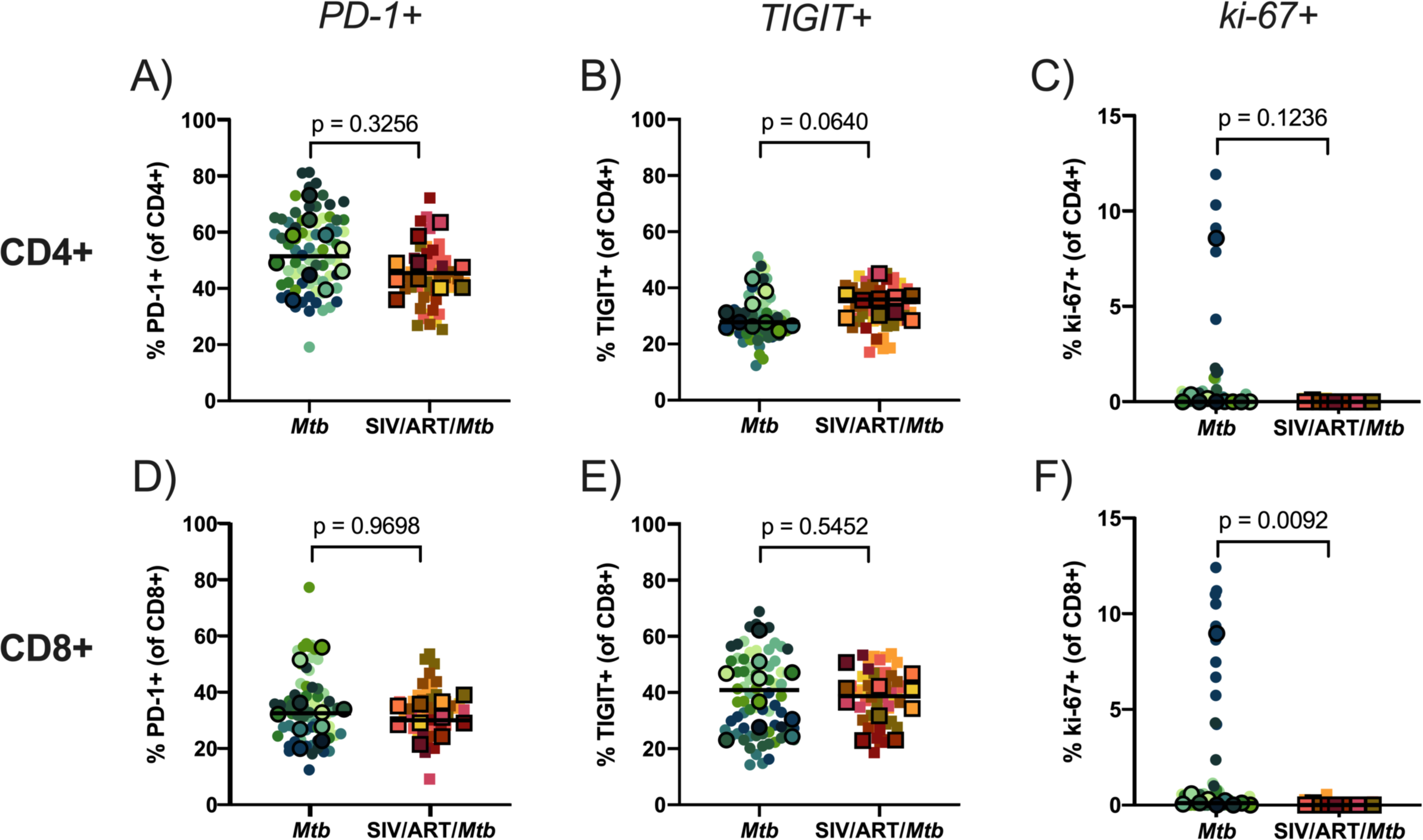
PD-1, TIGIT, and ki-67 frequency of CD4+ and CD8+ T cells isolated from granulomas. Frequencies of phenotype markers in granulomas relative to CD4+ or CD8+ gate. Outlined symbols indicate median per animal and unlined symbols indicate individual samples. Bars indicate group medians. Mann Whitney U tests of group medians were performed to determine significance. *P*-values are shown. A) PD-1+ CD4+ T cells. B) TIGIT+ CD4+ T cells. C) ki-67+ CD4+ T cells. D) PD-1+ CD8+ T cells. E) TIGIT+ CD8+ T cells. F) ki-67+ CD8+ T cells.

Our findings show that, while some differences were detected between SIV-infected, ART-treated juvenile macaques that are coinfected with Mtb, compared to macaques infected with just Mtb, the overall immune response and TB progression appear to be remarkably similar.

## Discussion

TB progression in HIV+ and HIV-naïve children is not well understood due to challenges in diagnosis, monitoring, and limited epidemiological data (1, 61). Studies from the pre-antibiotic era have noted that infants and young children are at greater risk of developing pulmonary TB and more severe, disseminated forms of TB (*e.g.*, miliary TB or TB meningitis) (9). This risk declines between the ages of 2 and 10 followed by an increased risk of pulmonary TB during puberty (9). HIV increases TB risk and results in more severe TB disease (2-4, 10, 11). It has yet to be thoroughly investigated how TB, both on its own and with a concurrent HIV infection, manifests in the presence of a developing immune system. Conventional and unconventional T cell compartments are in constant flux throughout childhood (13–16), highlighting the need to elucidate immune mechanisms behind TB pathogenesis where adult models may fall short. Only a handful of studies have modeled TB progression using young animals (45–47) and there are no studies that have modeled the impact of HIV coinfection on TB progression in juvenile macaques.

Here, we established a model of pediatric HIV/Mtb coinfection by infecting juvenile macaques with SIV, initiating ART 3 months later, and coinfecting with Mtb 3 months later. We compared TB progression in the SIV-infected, ART-treated juveniles with that in similarly aged animals infected with Mtb alone. We chose to implement ART in this study as ART coverage in children is improving worldwide (62) and is an increasingly likely real-world scenario. Prior to Mtb coinfection, the SIV-infected animals were aviremic, indicative of successful viral control by ART (Figure 1). Both groups exhibited similar progression of TB disease by PET/CT (Figure 4), pathology scores, Mtb dissemination and Mtb burden, with the exception of lower lung CFU in SIV-infected, ART-treated animals (Figures 5, 6; Figure S2). These results are in stark contrast with our previous study of adult macaques coinfected with SIV/Mtb in which all coinfected animals developed rapid dissemination of TB within the first 8 weeks of Mtb coinfection and extensive TB pathology (40). This difference is most likely due to the ART that was provided to the SIV-infected juveniles in the current study, consistent with the efficacy of ART reported in both adults (63, 64) and children (3–7, 65). We cannot rule out age as a potential contributor to the differences observed in TB progression in SIV-infected adult macaques (40) and the juveniles reported here since we did not coinfect SIV-infected, ART-naïve juveniles. However, clinical studies suggest that HIV-naïve children are slightly more susceptible, and not more resistant, to severe forms of TB than adults (8, 9). The striking degree to which TB disease in our SIV-infected juveniles paralleled that in SIV-naïve juveniles suggests strongly that ART may be responsible for constraining TB progression in SIV-infected juvenile macaques.

In humans, ART only partially restores immune function and resistance to Mtb in HIV+ subjects (63, 64). Thus, we characterized the T cell responses to Mtb in the SIV-infected, ART-treated animals relative to those in animals infected with Mtb alone. Since Mtb primarily establishes infection in the lung, we investigated T cell-mediated immunity in the airways throughout the study and in granulomas and lung tissue harvested 12 weeks after Mtb infection. We showed previously that SIV-related changes occur across T cell subsets in several lung compartments, including airways, granulomas, and lung tissue, in SIV-infected, ART-naïve adult macaques coinfected with Mtb (44). Adult macaques coinfected with SIV/Mtb exhibited fewer CD4+ T cells in blood and more CD8+ T cells in airways (44). Similarly, in the SIV-infected juvenile macaques here, we observed decreased CD4+ T cells and increased CD8+ T cells in the airways prior to ART initiation (Figure 2). However, following ART treatment, both T cell types returned to pre-SIV levels indicating a restoration of CD4+ and CD8+ T cell levels in the airways by ART. The unconventional T cell populations, Vγ9+ γδ T cells, MAIT and NKT cells, which are widely thought to play a role in early Mtb infection (29–33), varied in frequency between animals and there was no consistent change in these populations during SIV infection, ART, or Mtb coinfection. Following Mtb infection, no striking differences were observed between the groups in either conventional T cell subsets or unconventional T cells in the airways (Figure 2), aside from the elevated CD4+CD8+ T cells that is likely due to Mtb-mediated immune activation (48).

In addition to analyzing cells from the airway, we analyzed both lung tissue and individual granulomas. The latter is especially important since substantial heterogeneity can exist between granulomas, even from within the same animal (66). We did not observe differences in granuloma composition of T cell subsets, conventional or unconventional, nor were there any consistent differences in other tissues analyzed between SIV-naïve and SIV/ART-treated Mtb coinfected macaques (Figure 8 and Figure S3). IFNγ and TNF responses upon Mtb-specific stimulation of CD4+ and CD8+ T cells isolated from lungs were also similar between SIV-infected, ART-treated macaques coinfected with Mtb and macaques infected with Mtb alone (Figure 7). Both cytokines are associated with Mtb control (67, 68). Notably, we previously identified a defect in TNF production in SIV/Mtb coinfected adult macaques which may, at least in part, be responsible for the loss of control over Mtb in SIV-infected animals (44). A similar TNF defect in lymphocytes from lung tissue was not observed in the SIV-infected juvenile macaques studied here and may be attributed to the ART regimen. Mtb-specific responses in CD4+ and CD8+ T cells in blood from HIV/Mtb coinfected individuals have been shown to increase following ART, indicative of restoration of immune responses (69, 70). Consistent with other studies in human and NHP (26, 71), our data suggests that these similarities in IFNγ and TNF may be attributed to immune restoration from the ART regimen.

Chronic immune activation has been implicated in immune dysfunction associated with HIV/SIV infection (72, 73). PD-1 and TIGIT are inhibitory receptors that are critical in modulating immune activation and play an important role in Mtb control (74–77). Importantly, PD-1 and TIGIT expression can be upregulated on both exhausted, as well as activated, T cells. Previously, we found that frequencies of PD-1+ and TIGIT+ T cells are notably higher in lung tissue and granulomas from SIV-infected, ART-naïve adult macaques and appear to be activated, regardless of whether they are coinfected with Mtb (44). This is consistent with the hypothesis that this immune activation phenotype is due to chronic SIV infection and is maintained throughout SIV/Mtb coinfection (44, 73). Interestingly, in this study, there was no difference between the two groups of juvenile macaques in PD-1 or TIGIT expression on CD4+ and CD8+ T cells from either airways (Figure S1) or granulomas (Figure 10), aside from the decline in PD-1+ CD4+ T cells in airways following Mtb infection. Signaling pathways associated with inflammation and immune activation have been shown to decline during the first 6 months of ART in whole blood from HIV-1/Mtb coinfected individuals (78). Similarly, our data suggest that ART may have alleviated the chronic immune activation associated with uncontrolled viral replication.

We did observe transient changes in proliferation and cell trafficking in airways over the course of the SIV and Mtb infections. Proliferation of both CD4+ and CD8+ T cells spiked 4 weeks after SIV infection (Figure S1), which closely mirrored the spike in plasma viremia (Figure 1). Similar elevations in CD4+ and CD8+ T cell proliferation were previously observed in blood, lymph nodes, and gut shortly after SIV infection (49–51). This burst of T cell proliferation corresponds to elevations in T cell apoptosis (50) and likely indicates a compensatory response to restore CD4+ T cell levels depleted by rampant viral replication, while simultaneously boosting CD8+ T cells, well-known antiviral effectors cells. We observed an increase in the frequency of CCR6+CD4+ and CCR6+CD8+ T cells in both groups following Mtb infection. CCR6 is a chemokine receptor expressed on both dendritic cells and T cells (52). CCR6 expression on T cells promotes migration of Th1 and Th17 responses to sites of inflammation and it is associated with Mtb control (53). In our study, ART-treated SIV infection did not appear to affect the expression of CCR6 on CD4+ and CD8+ T cells in the airways during Mtb coinfection compared to animals infected with Mtb alone.

CCR5 is an important chemokine receptor for T cell migration during inflammation and is a co-receptor for HIV/SIV (22). Mtb-specific CD4 T cells upregulate CCR5 during latent TB infection and are preferentially depleted during HIV infection (79, 80). We observed a decline in the frequency of CCR5+ CD4+ T cells as well as a subtle, but significant, loss of absolute CCR5+ CD4+ T cells in the airways of SIV-infected, ART-treated macaques following Mtb coinfection (Figure 3). However, we did not observe a loss of CCR5+ CD4+ T cells in granulomas (Figure 9A). CCR5 is expressed at higher frequencies on T cells in airways compared to other compartments like blood (81). It is possible that the elevated expression of CCR5 on CD4 T cells in airways combined with Mtb coinfection created a target-rich environment for SIV, thereby resulting in their selective depletion (22). Interestingly, others have shown that CCR5+ CD4+ T cells are rapidly depleted in granulomas from untreated Mtb/SIV coinfected macaques (82). These data suggest that ART may have prevented depletion of CCR5+ CD4+ T cells in granulomas of our SIV-infected macaques. CCR5 is also be expressed on type 1 CD8+ T cells, which largely secrete IFNγ (55). SIV infection has been reported to cause significant depletion of CCR5+ CD8+ T cells at mucosal sites, such as jejunum (55). Others have reported CCR5 expression decline on circulating CD8+ T cells from HIV-infected progressors (83). We observed a significant decline in the frequency of CCR5+ CD8+ T cells and total CCR5+ CD8+ T cells in the airways after Mtb coinfection in SIV-infected, ART-treated juvenile macaques (Figure 3B, C). It is possible that CCR5 was downregulated on CD8+ T cells during Mtb coinfection, however, the decline of absolute CCR5+ CD8+ T cells suggests a loss of these cells. In granulomas, on the other hand, we did not observe a significant loss of CCR5+ CD8+ T cells (Figure 9B). Unlike CD4+ T cells, which have a clear mechanism of depletion via direct viral infection, the mechanism behind this tissue-dependent loss of CD8+ T cells during SIV infection is not well understood and warrants further investigation. Nevertheless, TB outcome did not differ between the two groups, indicating that the loss CCR5+ T cells in the airways did not exert a substantial effect on TB resistance.

In summary, we found that SIV-infected, ART-treated juvenile macaques control TB similarly to juvenile macaques infected with Mtb alone. Further, immune responses to Mtb do not appear to be substantially impaired in SIV-infected juvenile macaques on ART. One limitation of our study was that we did not have an ART-naïve group of SIV-infected juveniles with which to directly compare the effect of ART on TB pathogenesis in SIV-infected juvenile macaques. There is ample evidence in humans (69, 70, 78) that supports our hypothesis that ART may have restored anti-Mtb immunity and control of TB in SIV-infected juvenile macaques. ART previously was shown to reduce the risk of SIV-induced reactivation of latent TB infection (LTBI) in adult macaques (71), and ART may be an important tool to reduce TB reactivation in HIV+ people with LTBI (84). Our study is the first to suggest that early implementation of ART can restore the rate of TB progression in macaques already infected with SIV to that of SIV-naïve macaques, and the first time ART has been studied in a pediatric coinfection model. Whether later ART initiation would result in a similar host responses and rate of TB progression warrants further investigation. Although, early ART has been noted to have beneficial effects on CD8 T cells in HIV-infected children (85). Our data indicate that early ART initiation provides not only viral control but may also reduce TB severity and mortality in children living with HIV. Indeed, ART is highly beneficial in reducing mortality in children infected with HIV (86). Unfortunately, ART coverage worldwide in children still lags behind adults (62) and this study further justifies increasing ART coverage in this vulnerable population.

## Materials and Methods

### Animal studies

Juvenile (∼1-2 years of age, equivalent to children 4-8 years-old) Mauritian cynomolgus macaques (*Macaca fascicularis*) were obtained from Bioculture US (Immokalee, FL) (Table S1). MHC haplotype was determined by MiSeq sequencing and animals with the presence of at least one copy of the M1 MHC haplotype were selected for this study (87).

Animal protocols and procedures were approved by the University of Pittsburgh Institutional Animal Care and Use Committee (IACUC) which adheres to guidelines established in the Animal Welfare Act and the Guide for the Care and Use of Laboratory Animals, as well as the Weatherall Report (8th Edition). The University is fully accredited by AAALAC (accreditation number 000496), and its OLAW animal welfare assurance number is D16-00118. The IACUC reviewed and approved the study protocols 19014337 and 22010433, under Assurance Number A3187-01.

Animal welfare was monitored as described previously (44). In brief, all animals were checked at least twice daily to assess appetite, attitude, activity level, hydration status, etc. Following Mtb infection, the animals were monitored closely for clinical signs of TB (*e.g.*, weight loss, tachypnea, dyspnea, or coughing). Physical exams, including weights, were performed on a regular basis. Animals were sedated for all veterinary procedures (*e.g.*, blood draws) using ketamine or other approved drugs. Regular PET/CT imaging was conducted and has proven to be very useful for monitoring TB progression. Our veterinary technicians monitored animals especially closely for any signs of pain or distress. If any were noted, appropriate supportive care (*e.g.*, dietary supplementation and rehydration) and treatments (analgesics) were given. Any animal considered to have advanced disease or intractable pain from any cause, was deemed to have reached the humane endpoint, sedated with ketamine and humanely euthanized using sodium pentobarbital.

### SIV and Mtb infections of macaques

One group of juvenile macaques was infected intravenously with SIVmac239M (10,000 IU) (Figure S4). SIVmac239 is a molecularly barcoded virus stock generated from clonal SIVmac239 (88). A daily ART regimen of dolutegravir (DTG; 2.5 mg/mL, *s.c.*), tenofovir disoproxil fumarate (TDF; 5.1 mg/mL, *s.c.*), and emtricitabine (FTC; 40 mg/mL, *s.c.*) (89) was initiated three months after SIV infection and continued for the remainder of the study. DTG was kindly provided by ViiV Healthcare (Middlesex, UK) and TDF and FTC were kindly provided by Gilead (Foster City, CA).

For Mtb infection, SIV-naïve (n = 10) and SIV-infected (n = 10) juvenile animals were infected with a low dose (5 - 11 CFU) of Mtb Erdman via bronchoscopic instillation, as described previously (56), and followed for 12 weeks post Mtb infection (Figure 9).

### Clinical and microbiological monitoring

All animals were assessed twice daily for general health and monitored closely for clinical signs of TB (coughing, weight loss, tachypnea, dyspnea, etc.) following Mtb infection. Monthly gastric aspirates (GA) and bronchoalveolar lavage (BAL) samples were tested for Mtb growth. GA and BAL samples with culturable Mtb (+) or that were sterile (-) are indicated in Table S1. Blood was drawn at regular intervals to measure erythrocyte sedimentation rate and to provide peripheral blood mononuclear cells (PBMC) and plasma.

### PET/CT imaging and analysis

Radiolabeled 2-deoxy-2-(^18^F)fluoro-D-glucose (FDG) PET/CT was performed just prior to Mtb infection and then monthly after Mtb infection. Imaging was performed using a MultiScan LFER-150 PET/CT scanner (Mediso Medical Imaging Systems, Budapest, Hungary) housed within our BSL3 facility as previously described (90, 91). Co-registered PET/CT images were analyzed using OsiriX MD software (version 12.5.2, Pixmeo, Geneva, Switzerland) to enumerate granulomas and to calculate the total FDG avidity of the lungs, exclusive of lymph nodes, which is a quantitative measure of total inflammation in the lungs (90, 92). Thoracic lymphadenopathy and extrapulmonary dissemination of Mtb to the spleen and/or liver were also assessed qualitatively on these scans.

### Necropsy

Necropsies were performed as previously described (40, 44) 12 weeks after Mtb infection. One SIV-infected, ART-treated animal (35319) met humane endpoint criteria and necropsied 6 weeks after Mtb coinfection (Table S1). A final FDG PET/CT scan was performed no more than three days prior to necropsy to document disease progression and to guide collection of individual granulomas (56). Animals were heavily sedated with ketamine, maximally bled, and humanely euthanized using sodium pentobarbital (Beuthanasia, Schering-Plough, Kenilworth, NJ). Granulomas matched to the final PET/CT images were harvested along with other TB pathologies (e.g., consolidations and pneumonia), thoracic and extrathoracic lymph nodes, lung tissue, as well as portions of liver and spleen. Quantitative gross pathology scores were calculated and reflect overall TB disease burden for each animal (56). Tissue samples were divided and a portion was fixed in 10% neutral buffered formalin (NBF) for histopathology; the remainder was homogenized to a single-cell suspension as described previously (56). Serial dilutions of these homogenates were plated onto 7H11 agar, incubated at 37°C, 5% CO_2_ for three weeks, and colonies were enumerated. Samples yielding colonies were considered CFU+. Bacterial load in lungs, thoracic lymph nodes, liver, and spleen, as well as total thoracic CFU, were calculated as described previously (48). NBF-fixed tissue was embedded in paraffin, sectioned, and stained with hematoxylin and eosin for histopathologic examination.

### Flow cytometry of BAL and tissue

In general, cells collected from BAL and necropsy were stained following a similar protocol. Cells were counted and aliquoted at 1×10^6^ cells/well in a 96-well plate and stained immediately to ensure proper assessment of conventional and unconventional T cell subsets. Cells from lung lobes, lymph nodes, and spleen were stimulated for 14 hours total with either Mtb H37Rv whole cell lysate (20 μg/mL; BEI Resources) or *M. smegmatis* (MOI 5:1) at 37°C. In brief, stimulators were added and incubated for 2 hours, then brefeldin A (1 μg/mL) was added for the remainder of the stimulation time. Cells were reconstituted in 500 nM dasatinib in ELISpot media (RPMI 1640 + 10% human albumin + 1% glycine + 1% HEPES buffer) to improve tetramer staining. MR1 (5-OPRU; PE) and CD1d (PBS-57; APC) tetramers were added to wells and incubated for 30 minutes at room temperature (NIH Tetramer Core Facility, Atlanta, GA). Antibody for Vα7.2 (Table S2) was added, incubated for an additional 30 minutes at room temperature, and washed twice with PBS containing dasatinib. Cells were stained with a viability dye (Table S2) for 10 minutes at room temperature and washed with FACS containing dasatinib. Cells were stained with surface antibody cocktail (Table S2) for 20 minutes at 4°C, were fixed in 1% PFA, and permeabilized with BD Cytofix/Cytoperm™ (BD; Cat No. 554714) for 10 minutes at room temperature. Cells were stained intracellularly for 20 minutes at room temperature, washed, and analyzed immediately.

Flow cytometry of BAL and tissue samples was performed using a Cytek Aurora (BD). FCS files were analyzed using FlowJo software for Macintosh (version 10.1). Gating strategies for BAL and tissue data are shown in Figure S5 and S6, respectively. For most samples, we acquired 50,000 events in the lymphocyte gate. However, when this was not possible (i.e. for some small granulomas), we applied a cutoff threshold of CD3 events >100. Samples below that threshold were excluded from further analysis. Absolute CCR5+ CD4+ and CD8+ T cell counts were calculated by multiplying the frequency of CCR5+ or CCR5-CD4+ and CD8+ T cells from the live leukocyte gate with the hemocytometer count.

### Sequencing of barcode identifiers in Mtb

Mtb genomic DNA was isolated as previously described (58) and Mtb barcodes were sequenced as previously described (93). Only CFU+ samples were included in Mtb barcode analyses. Briefly, genomic DNA was isolated and samples were quantified and diluted to 10 ng/uL. Samples were then amplified twice using 2x Q5 Master Mix (New England BioLabs) and two unique primer sets, one to add a molecular counter to distinguish unique input templates, and one to add the Illumina TruSeq adapter sequences, were used. Primer sequences can be found in Table S3. Samples were then sequenced using a 2×150 MiSeq cartridge with v2 chemistry at a concentration of 4 pM and a 20% PhiX spike. A computational pipeline courtesy of Dr. Michael Chase and the Fortune lab was used to determine the number and proportion of unique barcode sequences in each sample from FASTQ files (93). Only barcodes present at a frequency of 1% or greater were included. All figures were generated using Prism 9 and Adobe Illustrator 2019.

### Statistics

For comparing longitudinal BAL data, linear mixed models with subject as a random variable were used to test treatment groups over time (Table S4). Fixed effect tests were used to assess whether there were differences among treatment groups or among time points. All time points were then compared with Tukey HSD (honestly significant difference) test using the Tukey-Kramer multiple comparison adjustment. Linear models were run in JMP® Pro (v.14.3.0). A two-way repeated measures ANOVA was used to test if there were differences in the number of barcodes between infection group (Mtb vs. SIV/ART/Mtb) and tissue type (granulomas vs. thoracic lymph nodes) and Bonferroni’s multiple comparison adjusted p-values were reported (Figure S2B).

For all other data, the Shapiro-Wilk normality test was used to check for normal distribution of data. Unpaired normally distributed data were analyzed using t tests, while unpaired non-normally distributed data were analyzed with the Mann-Whitney U test. Statistical tests were performed in Prism (version 8.2.1; GraphPad). All tests were two-sided, and statistical significance was designated at a *P* value of < 0.05. *P* values between 0.05 and 0.10 were considered trending.

### Sequence availability

All Mtb barcode sequences are available on the Sequence Read Archive (SRA) under accession number PRJNA900591.

## Acknowledgements

We would like to thank our collaborator at the University of Melbourne, Dr. Daniel Pellicci for his valuable discussions and technical input. We thank the veterinary and laboratory staffs of the TB Research Group at the University of Pittsburgh for their hard work. We are grateful to Dr. Philana Ling Lin, MD, MPH for necropsy assistance and to Dr. Edwin Klein, DVM for his expert review of the histopathology slides. We thank ViiV Healthcare for providing DTG and Gilead Sciences Inc for providing TDF and FTC. The MR1 tetramer technology was developed jointly by Dr. James McClusky, Dr. Jamie Rossjohn, and Dr. David Fairlie, and the material was produced by the NIH Tetramer Core Facility as permitted by the University of Melbourne (94). We thank the NIH Tetramer Core Facility (contract number 75N93020D00005) for providing MR1 and CD1d tetramers. This work was supported by the National Institutes of Health award AI142662.

## Supplementary Materials

**Table S1. Juvenile macaques *Mtb* outcome data**

**Table S2. List of antibodies used for flow cytometry.**

**Table S3. List of primers used for uniquely tagged Mtb.**

**Table S4. Statistics of BAL T cell composition.** A mixed statistical model was performed to determine significance with fixed effects (time and group) and random effects (individual animals). Tukey Pairwise comparisons were performed between time points.

**Figure S1. Phenotype of CD3+CD4+ and CD3+CD8+ T cells in airways over time.** Proliferation (ki-67), activation/exhaustion (PD-1 & TIGIT), and chemokine receptors (CXCR3 & CCR6) were measured on CD4+ and CD8+ T cells by flow cytometry. Individual animals indicated by symbols. SIV-infected, ART-treated Mtb coinfected animals are indicated by orange lines and animals infected with Mtb alone are indicated by teal lines. A mixed statistical model was performed to determine significance with fixed effects (time and group) and random effects (individual animals). Bars with asterisks indicate significance between time points; # 0.05 < p < 0.1, * p < 0.05, ** p < 0.01, *** p < 0.001 and **** p < 0.0001.

**Figure S2. Bacterial dissemination in macaques infected with Mtb alone or SIV-infected, ART-treated Mtb coinfected macaques.** A) Total number of uniquely tagged Mtb lineages found in each animal. Significance was determined using Mann Whitney U tests. Each dot indicates an individual animal. B) Number of barcodes identified within granulomas or thoracic lymph nodes. A two-way repeated measures ANOVA with Bonferroni’s multiple comparison was used to determine differences in the number of barcodes between infection group and tissue type. Each dot indicates an individual animal.

**Figure S3. T cell subsets in lung tissue, thoracic lymph nodes, extrathoracic lymph nodes, spleen, and PBMC.** Frequencies of T cell subsets relative to CD3+ gate. Outlined symbols indicate median per animal and unlined symbols indicate individual samples. Bars indicate group medians. Mann Whitney U tests of group medians were performed to determine significance. *P*- values are shown. A) CD4+ T cells. B) CD8+ T cells. C) CD4+CD8+ T cells. D) CD4-CD8-T cells. E) Vγ9+ γδ T cells. F) CD1d tet+ NKT cells. G) MR1 tet+ MAIT cells.

**Figure S4. Timeline of juvenile SIV-infected, ART-treated Mtb co-infection study.**

**Figure S5. Representative gating strategy of BAL.** A) BAL cells were stained with antibodies in Table S2 according to the methods. Shown is a representative gating schematic for unconventional T cells (Vγ9+ γδ T cells, CD3+Vγ9+; MAIT cells, CD3+ MR1 5-OP-RU tetramer+; NKT cells, CD3+ CD1d PBS-57 tetramer+) and conventional T cells (CD3+CD8+ and CD3+CD4+ cells after exclusion of markers from unconventional T cells). B) T cell subset populations (if event count > 50-100) were gated for the following markers: ki-67, PD-1, TIGIT, CCR5, CCR6, and CXCR3.

**Figure S6. Representative gating strategy of necropsy tissues (granulomas, lung, thoracic and extrathoracic lymph nodes, spleen, and PBMC).** A) Tissue samples collected at necropsy were stained with antibodies in Table S2 according to the methods. Shown is a representative gating schematic for unconventional T cells (NKT cells, CD3+ CD1d PBS-57 tetramer+; MAIT cells, CD3+ MR1 5-OP-RU tetramer+; Vγ9+ γδ T cells, CD3+Vγ9+) and conventional T cells (CD3+CD8+ and CD3+CD4+ cells after exclusion of markers from unconventional T cells). B) In instances where the parent population of cells was greater than 50-100 events, unconventional and conventional T cells were then gated for expression of memory markers (CD28/CD95) as well as CCR5, CCR7, PD-1, ki-67, TIGIT, Granzyme B, TNF, and IFNγ.

## References

1. World Health Organization. 2020. Global tuberculosis report 2020. Geneva: World Health Organization. https://www.who.int/publications/i/item/9789240013131

2. Palme IB, Gudetta B, Bruchfeld J, Muhe L, Giesecke J. 2002. Impact of human immunodeficiency virus 1 infection on clinical presentation, treatment outcome and survival in a cohort of Ethiopian children with tuberculosis. Pediatr Infect Dis J 21:1053–61.

3. Dodd PJ, Prendergast AJ, Beecroft C, Kampmann B, Seddon JA. 2017. The impact of HIV and antiretroviral therapy on TB risk in children: a systematic review and meta-analysis. Thorax 72:559–575.

4. Braitstein P, Nyandiko W, Vreeman R, Wools-Kaloustian K, Sang E, Musick B, Sidle J, Yiannoutsos C, Ayaya S, Carter EJ. 2009. The clinical burden of tuberculosis among human immunodeficiency virus-infected children in Western Kenya and the impact of combination antiretroviral treatment. Pediatr Infect Dis J 28:626–32.

5. Mu W, Zhao Y, Sun X, Ma Y, Yu L, Liu X, Zhao D, Dou Z, Fang H, Zhang F. 2014. Incidence and associated factors of pulmonary tuberculosis in HIV-infected children after highly active antiretroviral therapy (HAART) in China: a retrospective study. AIDS Care 26:1127–35.

6. Martinson NA, Moultrie H, van Niekerk R, Barry G, Coovadia A, Cotton M, Violari A, Gray GE, Chaisson RE, McIntyre JA, Meyers T. 2009. HAART and risk of tuberculosis in HIV-infected South African children: a multi-site retrospective cohort. Int J Tuberc Lung Dis 13:862–7.

7. Jensen J, Alvaro-Meca A, Micheloud D, Diaz A, Resino S. 2012. Reduction in mycobacterial disease among HIV-infected children in the highly active antiretroviral therapy era (1997-2008). Pediatr Infect Dis J 31:278–83.

8. Jenkins HE, Yuen CM, Rodriguez CA, Nathavitharana RR, McLaughlin MM, Donald P, Marais BJ, Becerra MC. 2017. Mortality in children diagnosed with tuberculosis: a systematic review and meta-analysis. Lancet Infect Dis 17:285–295.

9. Marais BJ, Gie RP, Schaaf HS, Hesseling AC, Obihara CC, Starke JJ, Enarson DA, Donald PR, Beyers N. 2004. The natural history of childhood intra-thoracic tuberculosis: a critical review of literature from the pre-chemotherapy era. Int J Tuberc Lung Dis 8:392–402.

10. Schaaf HS, Marais BJ, Whitelaw A, Hesseling AC, Eley B, Hussey GD, Donald PR. 2007. Culture-confirmed childhood tuberculosis in Cape Town, South Africa: a review of 596 cases. BMC Infect Dis 7:140.

11. Madhi SA, Huebner RE, Doedens L, Aduc T, Wesley D, Cooper PA. 2000. HIV-1 co-infection in children hospitalised with tuberculosis in South Africa. Int J Tuberc Lung Dis 4:448–54.

12. Simon AK, Hollander GA, McMichael A. 2015. Evolution of the immune system in humans from infancy to old age. Proc Biol Sci 282:20143085.

13. Trück J, van der Burg M. 2020. Development of adaptive immune cells and receptor repertoires from infancy to adulthood. Current Opinion in Systems Biology 24:51–55.

14. Kumar BV, Connors TJ, Farber DL. 2018. Human T Cell Development, Localization, and Function throughout Life. Immunity 48:202–213.

15. Shearer WT, Rosenblatt HM, Gelman RS, Oyomopito R, Plaeger S, Stiehm ER, Wara DW, Douglas SD, Luzuriaga K, McFarland EJ, Yogev R, Rathore MH, Levy W, Graham BL, Spector SA. 2003. Lymphocyte subsets in healthy children from birth through 18 years of age: the Pediatric AIDS Clinical Trials Group P1009 study. J Allergy Clin Immunol 112:973–80.

16. Jalali S, Harpur CM, Piers AT, Auladell M, Perriman L, Li S, An K, Anderson J, Berzins SP, Licciardi PV, Ashhurst TM, Konstantinov IE, Pellicci DG. 2022. A high-dimensional cytometry atlas of peripheral blood over the human life span. Immunol Cell Biol 100:805–821.

17. Janković V, Messaoudi I, Nikolich-Zugich J. 2003. Phenotypic and functional T-cell aging in rhesus macaques (Macaca mulatta): differential behavior of CD4 and CD8 subsets. Blood 102:3244–51.

18. Pitcher CJ, Hagen SI, Walker JM, Lum R, Mitchell BL, Maino VC, Axthelm MK, Picker LJ. 2002. Development and homeostasis of T cell memory in rhesus macaque. J Immunol 168:29–43.

19. Serbina NV, Lazarevic V, Flynn JL. 2001. CD4(+) T cells are required for the development of cytotoxic CD8(+) T cells during Mycobacterium tuberculosis infection. J Immunol 167:6991–7000.

20. van Pinxteren LA, Cassidy JP, Smedegaard BH, Agger EM, Andersen P. 2000. Control of latent Mycobacterium tuberculosis infection is dependent on CD8 T cells. Eur J Immunol 30:3689–98.

21. Baker CA, Clark R, Ventura F, Jones NG, Guzman D, Bangsberg DR, Cao H. 2007. Peripheral CD4 loss of regulatory T cells is associated with persistent viraemia in chronic HIV infection. Clin Exp Immunol 147:533–9.

22. Okoye AA, Picker LJ. 2013. CD4(+) T-cell depletion in HIV infection: mechanisms of immunological failure. Immunol Rev 254:54–64.

23. Hoffmann M, Pantazis N, Martin GE, Hickling S, Hurst J, Meyerowitz J, Willberg CB, Robinson N, Brown H, Fisher M, Kinloch S, Babiker A, Weber J, Nwokolo N, Fox J, Fidler S, Phillips R, Frater J, Spartac, Investigators C. 2016. Exhaustion of Activated CD8 T Cells Predicts Disease Progression in Primary HIV-1 Infection. PLoS Pathog 12:e1005661.

24. Fenwick C, Joo V, Jacquier P, Noto A, Banga R, Perreau M, Pantaleo G. 2019. T-cell exhaustion in HIV infection. Immunol Rev 292:149–163.

25. Rutishauser RL, Hartogensis W, Deguit CD, Krone M, Hoh R, Hecht FM, Pilcher CD, Bacchetti P, Deeks SG, Hunt PW, McCune JM. 2017. Early and Delayed Antiretroviral Therapy Results in Comparable Reductions in CD8(+) T Cell Exhaustion Marker Expression. AIDS Res Hum Retroviruses 33:658–667.

26. Jensen SS, Fomsgaard A, Larsen TK, Tingstedt JL, Gerstoft J, Kronborg G, Pedersen C, Karlsson I. 2015. Initiation of Antiretroviral Therapy (ART) at Different Stages of HIV-1 Disease Is Not Associated with the Proportion of Exhausted CD8+ T Cells. PLoS One 10:e0139573.

27. Gras L, May M, Ryder LP, Trickey A, Helleberg M, Obel N, Thiebaut R, Guest J, Gill J, Crane H, Dias Lima V, d’Arminio Monforte A, Sterling TR, Miro J, Moreno S, Stephan C, Smith C, Tate J, Shepherd L, Saag M, Rieger A, Gillor D, Cavassini M, Montero M, Ingle SM, Reiss P, Costagliola D, Wit F, Sterne J, de Wolf F, Geskus R. 2019. Determinants of Restoration of CD4 and CD8 Cell Counts and Their Ratio in HIV-1-Positive Individuals With Sustained Virological Suppression on Antiretroviral Therapy. J Acquir Immune Defic Syndr 80:292–300.

28. Alvarez P, Mwamzuka M, Marshed F, Kravietz A, Ilmet T, Ahmed A, Borkowsky W, Khaitan A. 2017. Immune activation despite preserved CD4 T cells in perinatally HIV-infected children and adolescents. PLoS One 12:e0190332.

29. Shao L, Zhang W, Zhang S, Chen CY, Jiang W, Xu Y, Meng C, Weng X, Chen ZW. 2008. Potent immune responses of Ag-specific Vgamma2Vdelta2+ T cells and CD8+ T cells associated with latent stage of Mycobacterium tuberculosis coinfection in HIV-1-infected humans. Aids 22:2241–50.

30. Shen L, Frencher J, Huang D, Wang W, Yang E, Chen CY, Zhang Z, Wang R, Qaqish A, Larsen MH, Shen H, Porcelli SA, Jacobs WR, Jr., Chen ZW. 2019. Immunization of Vγ2Vδ2 T cells programs sustained effector memory responses that control tuberculosis in nonhuman primates. Proc Natl Acad Sci U S A 116:6371–6378.

31. Gold MC, Cerri S, Smyk-Pearson S, Cansler ME, Vogt TM, Delepine J, Winata E, Swarbrick GM, Chua WJ, Yu YY, Lantz O, Cook MS, Null MD, Jacoby DB, Harriff MJ, Lewinsohn DA, Hansen TH, Lewinsohn DM. 2010. Human mucosal associated invariant T cells detect bacterially infected cells. PLoS Biol 8:e1000407.

32. Ulrichs T, Moody DB, Grant E, Kaufmann SH, Porcelli SA. 2003. T-cell responses to CD1-presented lipid antigens in humans with Mycobacterium tuberculosis infection. Infect Immun 71:3076–87.

33. Sada-Ovalle I, Chiba A, Gonzales A, Brenner MB, Behar SM. 2008. Innate invariant NKT cells recognize Mycobacterium tuberculosis-infected macrophages, produce interferon-gamma, and kill intracellular bacteria. PLoS Pathog 4:e1000239.

34. Juno JA, Eriksson EM. 2019. γδ T-cell responses during HIV infection and antiretroviral therapy. Clin Transl Immunology 8:e01069.

35. Leeansyah E, Ganesh A, Quigley MF, Sonnerborg A, Andersson J, Hunt PW, Somsouk M, Deeks SG, Martin JN, Moll M, Shacklett BL, Sandberg JK. 2013. Activation, exhaustion, and persistent decline of the antimicrobial MR1-restricted MAIT-cell population in chronic HIV-1 infection. Blood 121:1124–35.

36. Fernandez CS, Kelleher AD, Finlayson R, Godfrey DI, Kent SJ. 2014. NKT cell depletion in humans during early HIV infection. Immunol Cell Biol 92:578–90.

37. Capuano SV, 3rd, Croix DA, Pawar S, Zinovik A, Myers A, Lin PL, Bissel S, Fuhrman C, Klein E, Flynn JL. 2003. Experimental Mycobacterium tuberculosis infection of cynomolgus macaques closely resembles the various manifestations of human M. tuberculosis infection. Infect Immun 71:5831–44.

38. Scanga CA, Flynn JL. 2014. Modeling tuberculosis in nonhuman primates. Cold Spring Harb Perspect Med 4:a018564.

39. Schmitz JE, Korioth-Schmitz B. 2013. Immunopathogenesis of simian immunodeficiency virus infection in nonhuman primates. Curr Opin HIV AIDS 8:273–9.

40. Rodgers MA, Ameel C, Ellis-Connell AL, Balgeman AJ, Maiello P, Barry GL, Friedrich TC, Klein E, O’Connor SL, Scanga CA. 2018. Preexisting Simian Immunodeficiency Virus Infection Increases Susceptibility to Tuberculosis in Mauritian Cynomolgus Macaques. Infect Immun 86.

41. Kerkhoff AD, Barr DA, Schutz C, Burton R, Nicol MP, Lawn SD, Meintjes G. 2017. Disseminated tuberculosis among hospitalised HIV patients in South Africa: a common condition that can be rapidly diagnosed using urine-based assays. Sci Rep 7:10931.

42. Rugină S, Dumitru IM, Resul G, Cernat RC, Petcu AE. 2014. Disseminated tuberculosis in HIV-infected patients from the Regional HIV/AIDS Center Constanţa, Romania. Germs 4:16–21.

43. Schutz C, Barr D, Andrade BB, Shey M, Ward A, Janssen S, Burton R, Wilkinson KA, Sossen B, Fukutani KF, Nicol M, Maartens G, Wilkinson RJ, Meintjes G. 2019. Clinical, microbiologic, and immunologic determinants of mortality in hospitalized patients with HIV-associated tuberculosis: A prospective cohort study. PLOS Medicine 16:e1002840.

44. Larson EC, Ellis-Connell A, Rodgers MA, Balgeman AJ, Moriarty RV, Ameel CL, Baranowski TM, Tomko JA, Causgrove CM, Maiello P, O’Connor SL, Scanga CA. 2021. Pre-existing Simian Immunodeficiency Virus Infection Increases Expression of T Cell Markers Associated with Activation during Early Mycobacterium tuberculosis Coinfection and Impairs TNF Responses in Granulomas. J Immunol 207:175–188.

45. Cepeda M, Salas M, Folwarczny J, Leandro AC, Hodara VL, de la Garza MA, Dick EJ, Jr., Owston M, Armitige LY, Gauduin MC. 2013. Establishment of a neonatal rhesus macaque model to study Mycobacterium tuberculosis infection. Tuberculosis (Edinb) 93 Suppl:S51–9.

46. Chen CY, Huang D, Wang RC, Shen L, Zeng G, Yao S, Shen Y, Halliday L, Fortman J, McAllister M, Estep J, Hunt R, Vasconcelos D, Du G, Porcelli SA, Larsen MH, Jacobs WR, Jr., Haynes BF, Letvin NL, Chen ZW. 2009. A Critical Role for CD8 T Cells in a Nonhuman Primate Model of Tuberculosis. PLOS Pathogens 5:e1000392.

47. Qiu L, Huang D, Chen CY, Wang R, Shen L, Shen Y, Hunt R, Estep J, Haynes BF, Jacobs WR, Jr., Letvin N, Du G, Chen ZW. 2008. Severe tuberculosis induces unbalanced up-regulation of gene networks and overexpression of IL-22, MIP-1alpha, CCL27, IP-10, CCR4, CCR5, CXCR3, PD1, PDL2, IL-3, IFN-beta, TIM1, and TLR2 but low antigen-specific cellular responses. J Infect Dis 198:1514–9.

48. Diedrich CR, Gideon HP, Rutledge T, Baranowski TM, Maiello P, Myers AJ, Lin PL. 2019. CD4CD8 Double Positive T cell responses during Mycobacterium tuberculosis infection in cynomolgus macaques. J Med Primatol 48:82–89.

49. Benlhassan-Chahour K, Penit C, Dioszeghy V, Vasseur F, Janvier G, Rivière Y, Dereuddre-Bosquet N, Dormont D, Le Grand R, Vaslin B. 2003. Kinetics of lymphocyte proliferation during primary immune response in macaques infected with pathogenic simian immunodeficiency virus SIVmac251: preliminary report of the effect of early antiviral therapy. J Virol 77:12479–93.

50. Pandrea I, Gaufin T, Gautam R, Kristoff J, Mandell D, Montefiori D, Keele BF, Ribeiro RM, Veazey RS, Apetrei C. 2011. Functional cure of SIVagm infection in rhesus macaques results in complete recovery of CD4+ T cells and is reverted by CD8+ cell depletion. PLoS Pathog 7:e1002170.

51. Koopman G, Mortier D, Hofman S, Koutsoukos M, Bogers W, Wahren B, Voss G, Heeney JL. 2009. Acute-phase CD4+ T-cell proliferation and CD152 upregulation predict set-point virus replication in vaccinated simian-human immunodeficiency virus strain 89.6p-infected macaques. J Gen Virol 90:915–926.

52. Ito T, Carson WFt, Cavassani KA, Connett JM, Kunkel SL. 2011. CCR6 as a mediator of immunity in the lung and gut. Exp Cell Res 317:613–9.

53. Shanmugasundaram U, Bucsan AN, Ganatra SR, Ibegbu C, Quezada M, Blair RV, Alvarez X, Velu V, Kaushal D, Rengarajan J. 2020. Pulmonary Mycobacterium tuberculosis control associates with CXCR3- and CCR6-expressing antigen-specific Th1 and Th17 cell recruitment. JCI Insight 5.

54. Ahsan MH, Gill AF, Lackner AA, Veazey RS. 2012. Acute and chronic T cell dynamics in the livers of simian immunodeficiency virus-infected macaques. J Virol 86:5244–52.

55. Wang X, Russell-Lodrigue KE, Ratterree MS, Veazey RS, Xu H. 2019. Chemokine receptor CCR5 correlates with functional CD8(+) T cells in SIV-infected macaques and the potential effects of maraviroc on T-cell activation. Faseb j 33:8905–8912.

56. Maiello P, DiFazio RM, Cadena AM, Rodgers MA, Lin PL, Scanga CA, Flynn JL. 2018. Rhesus Macaques Are More Susceptible to Progressive Tuberculosis than Cynomolgus Macaques: a Quantitative Comparison. Infect Immun 86.

57. Lin PL, Ford CB, Coleman MT, Myers AJ, Gawande R, Ioerger T, Sacchettini J, Fortune SM, Flynn JL. 2014. Sterilization of granulomas is common in active and latent tuberculosis despite within-host variability in bacterial killing. Nat Med 20:75–9.

58. Martin CJ, Cadena AM, Leung VW, Lin PL, Maiello P, Hicks N, Chase MR, Flynn JL, Fortune SM. 2017. Digitally Barcoding Mycobacterium tuberculosis Reveals In Vivo Infection Dynamics in the Macaque Model of Tuberculosis. MBio 8.

59. de Noronha AL, Báfica A, Nogueira L, Barral A, Barral-Netto M. 2008. Lung granulomas from Mycobacterium tuberculosis/HIV-1 co-infected patients display decreased in situ TNF production. Pathol Res Pract 204:155–61.

60. Diedrich CR, O’Hern J, Wilkinson RJ. 2016. HIV-1 and the Mycobacterium tuberculosis granuloma: A systematic review and meta-analysis. Tuberculosis (Edinb) 98:62–76.

61. Thomas TA. 2017. Tuberculosis in Children. Pediatr Clin North Am 64:893–909.

62. UNAIDS. 2021. UNAIDS data 2021. https://www.unaids.org/en/resources/documents/2021/2021_unaids_data

63. Lawn SD, Myer L, Edwards D, Bekker LG, Wood R. 2009. Short-term and long-term risk of tuberculosis associated with CD4 cell recovery during antiretroviral therapy in South Africa. AIDS 23:1717–25.

64. Gupta A, Wood R, Kaplan R, Bekker LG, Lawn SD. 2012. Tuberculosis incidence rates during 8 years of follow-up of an antiretroviral treatment cohort in South Africa: comparison with rates in the community. PLoS One 7:e34156.

65. Anigilaje EA, Aderibigbe SA, Adeoti AO, Nweke NO. 2016. Tuberculosis, before and after Antiretroviral Therapy among HIV-Infected Children in Nigeria: What Are the Risk Factors? PLoS One 11:e0156177.

66. Gideon HP, Phuah J, Myers AJ, Bryson BD, Rodgers MA, Coleman MT, Maiello P, Rutledge T, Marino S, Fortune SM, Kirschner DE, Lin PL, Flynn JL. 2015. Variability in tuberculosis granuloma T cell responses exists, but a balance of pro- and anti-inflammatory cytokines is associated with sterilization. PLoS Pathog 11:e1004603.

67. Green AM, Difazio R, Flynn JL. 2013. IFN-γ from CD4 T cells is essential for host survival and enhances CD8 T cell function during Mycobacterium tuberculosis infection. J Immunol 190:270–7.

68. Allie N, Grivennikov SI, Keeton R, Hsu NJ, Bourigault ML, Court N, Fremond C, Yeremeev V, Shebzukhov Y, Ryffel B, Nedospasov SA, Quesniaux VF, Jacobs M. 2013. Prominent role for T cell-derived tumour necrosis factor for sustained control of Mycobacterium tuberculosis infection. Sci Rep 3:1809.

69. Chiacchio T, Petruccioli E, Vanini V, Cuzzi G, La Manna MP, Orlando V, Pinnetti C, Sampaolesi A, Antinori A, Caccamo N, Goletti D. 2018. Impact of antiretroviral and tuberculosis therapies on CD4(+) and CD8(+) HIV/M. tuberculosis-specific T-cell in co-infected subjects. Immunol Lett 198:33–43.

70. Riou C, Jhilmeet N, Rangaka MX, Wilkinson RJ, Wilkinson KA. 2020. Tuberculosis Antigen-Specific T-Cell Responses During the First 6 Months of Antiretroviral Treatment. J Infect Dis 221:162–167.

71. Sharan R, Ganatra SR, Bucsan AN, Cole J, Singh DK, Alvarez X, Gough M, Alvarez C, Blakley A, Ferdin J, Thippeshappa R, Singh B, Escobedo R, Shivanna V, Dick EJ, Jr., Hall-Ursone S, Khader SA, Mehra S, Rengarajan J, Kaushal D. 2022. Antiretroviral therapy timing impacts latent tuberculosis infection reactivation in a Mycobacterium tuberculosis/SIV coinfection model. J Clin Invest 132.

72. Sokoya T, Steel HC, Nieuwoudt M, Rossouw TM. 2017. HIV as a Cause of Immune Activation and Immunosenescence. Mediators of Inflammation 2017:6825493.

73. Sharan R, Bucşan AN, Ganatra S, Paiardini M, Mohan M, Mehra S, Khader SA, Kaushal D. 2020. Chronic Immune Activation in TB/HIV Co-infection. Trends Microbiol 28:619–632.

74. Pollock KM, Montamat-Sicotte DJ, Grass L, Cooke GS, Kapembwa MS, Kon OM, Sampson RD, Taylor GP, Lalvani A. 2016. PD-1 Expression and Cytokine Secretion Profiles of Mycobacterium tuberculosis-Specific CD4+ T-Cell Subsets; Potential Correlates of Containment in HIV-TB Co-Infection. PLoS One 11:e0146905.

75. Kauffman KD, Sakai S, Lora NE, Namasivayam S, Baker PJ, Kamenyeva O, Foreman TW, Nelson CE, Oliveira-de-Souza D, Vinhaes CL, Yaniv Z, Lindestam Arleham CS, Sette A, Freeman GJ, Moore R, Sher A, Mayer-Barber KD, Andrade BB, Kabat J, Via LE, Barber DL. 2021. PD-1 blockade exacerbates Mycobacterium tuberculosis infection in rhesus macaques. Sci Immunol 6.

76. Day CL, Abrahams DA, Bunjun R, Stone L, de Kock M, Walzl G, Wilkinson RJ, Burgers WA, Hanekom WA. 2018. PD-1 Expression on Mycobacterium tuberculosis-Specific CD4 T Cells Is Associated With Bacterial Load in Human Tuberculosis. Front Immunol 9:1995.

77. Chew GM, Fujita T, Webb GM, Burwitz BJ, Wu HL, Reed JS, Hammond KB, Clayton KL, Ishii N, Abdel-Mohsen M, Liegler T, Mitchell BI, Hecht FM, Ostrowski M, Shikuma CM, Hansen SG, Maurer M, Korman AJ, Deeks SG, Sacha JB, Ndhlovu LC. 2016. TIGIT Marks Exhausted T Cells, Correlates with Disease Progression, and Serves as a Target for Immune Restoration in HIV and SIV Infection. PLoS Pathog 12:e1005349.

78. Wilkinson KA, Schneider-Luftman D, Lai R, Barrington C, Jhilmeet N, Lowe DM, Kelly G, Wilkinson RJ. 2021. Antiretroviral Treatment-Induced Decrease in Immune Activation Contributes to Reduced Susceptibility to Tuberculosis in HIV-1/Mtb Co-infected Persons. Front Immunol 12:645446.

79. Geldmacher C, Schuetz A, Ngwenyama N, Casazza JP, Sanga E, Saathoff E, Boehme C, Geis S, Maboko L, Singh M, Minja F, Meyerhans A, Koup RA, Hoelscher M. 2008. Early depletion of Mycobacterium tuberculosis-specific T helper 1 cell responses after HIV-1 infection. J Infect Dis 198:1590–8.

80. Geldmacher C, Ngwenyama N, Schuetz A, Petrovas C, Reither K, Heeregrave EJ, Casazza JP, Ambrozak DR, Louder M, Ampofo W, Pollakis G, Hill B, Sanga E, Saathoff E, Maboko L, Roederer M, Paxton WA, Hoelscher M, Koup RA. 2010. Preferential infection and depletion of Mycobacterium tuberculosis-specific CD4 T cells after HIV-1 infection. J Exp Med 207:2869–81.

81. Santucci MB, Bocchino M, Garg SK, Marruchella A, Colizzi V, Saltini C, Fraziano M. 2004. Expansion of CCR5+ CD4+ T-lymphocytes in the course of active pulmonary tuberculosis. Eur Respir J 24:638–43.

82. Foreman TW, Nelson CE, Kauffman KD, Lora NE, Vinhaes CL, Dorosky DE, Sakai S, Gomez F, Fleegle JD, Parham M, Perera SR, Lindestam Arlehamn CS, Sette A, Brenchley JM, Queiroz ATL, Andrade BB, Kabat J, Via LE, Barber DL. 2022. CD4 T cells are rapidly depleted from tuberculosis granulomas following acute SIV co-infection. Cell Rep 39:110896.

83. Meijerink H, Indrati AR, van Crevel R, Joosten I, Koenen H, van der Ven AJAM. 2014. The number of CCR5 expressing CD4+ T lymphocytes is lower in HIV-infected long-term non-progressors with viral control compared to normal progressors: a cross-sectional study. BMC Infectious Diseases 14:683.

84. Sterling TR, Lin PL. 2020. Treatment of latent M. tuberculosis infection and use of antiretroviral therapy to prevent tuberculosis. J Clin Invest 130:5102–5104.

85. Frange P, Montange T, Le Chenadec J, Batalie D, Fert I, Dollfus C, Faye A, Blanche S, Chacé A, Fourcade C, Hau I, Levine M, Mahlaoui N, Marcou V, Tabone M-D, Veber F, Hoctin A, Wack T, Avettand-Fenoël V, Warszawski J, Buseyne F. 2021. Impact of Early Versus Late Antiretroviral Treatment Initiation on Naive T Lymphocytes in HIV-1-Infected Children and Adolescents –The-ANRS-EP59-CLEAC Study. Frontiers in Immunology 12.

86. Violari A, Cotton MF, Gibb DM, Babiker AG, Steyn J, Madhi SA, Jean-Philippe P, McIntyre JA. 2008. Early antiretroviral therapy and mortality among HIV-infected infants. N Engl J Med 359:2233–44.

87. Budde ML, Wiseman RW, Karl JA, Hanczaruk B, Simen BB, O’Connor DH. 2010. Characterization of Mauritian cynomolgus macaque major histocompatibility complex class I haplotypes by high-resolution pyrosequencing. Immunogenetics 62:773–80.

88. Fennessey CM, Pinkevych M, Immonen TT, Reynaldi A, Venturi V, Nadella P, Reid C, Newman L, Lipkey L, Oswald K, Bosche WJ, Trivett MT, Ohlen C, Ott DE, Estes JD, Del Prete GQ, Lifson JD, Davenport MP, Keele BF. 2017. Genetically-barcoded SIV facilitates enumeration of rebound variants and estimation of reactivation rates in nonhuman primates following interruption of suppressive antiretroviral therapy. PLoS Pathog 13:e1006359.

89. Borducchi EN, Cabral C, Stephenson KE, Liu J, Abbink P, Ng’ang’a D, Nkolola JP, Brinkman AL, Peter L, Lee BC, Jimenez J, Jetton D, Mondesir J, Mojta S, Chandrashekar A, Molloy K, Alter G, Gerold JM, Hill AL, Lewis MG, Pau MG, Schuitemaker H, Hesselgesser J, Geleziunas R, Kim JH, Robb ML, Michael NL, Barouch DH. 2016. Ad26/MVA therapeutic vaccination with TLR7 stimulation in SIV-infected rhesus monkeys. Nature 540:284–287.

90. Hartman AL, Nambulli S, McMillen CM, White AG, Tilston-Lunel NL, Albe JR, Cottle E, Dunn MD, Frye LJ, Gilliland TH, Olsen EL, O’Malley KJ, Schwarz MM, Tomko JA, Walker RC, Xia M, Hartman MS, Klein E, Scanga CA, Flynn JL, Klimstra WB, McElroy AK, Reed DS, Duprex WP. 2020. SARS-CoV-2 infection of African green monkeys results in mild respiratory disease discernible by PET/CT imaging and shedding of infectious virus from both respiratory and gastrointestinal tracts. PLoS Pathog 16:e1008903.

91. Sarnyai Z, Nagy K, Patay G, Molnár M, Rosenqvist G, Tóth M, Takano A, Gulyás B, Major P, Halldin C, Varrone A. 2019. Performance Evaluation of a High-Resolution Nonhuman Primate PET/CT System. J Nucl Med 60:1818–1824.

92. White AG, Maiello P, Coleman MT, Tomko JA, Frye LJ, Scanga CA, Lin PL, Flynn JL. 2017. Analysis of 18FDG PET/CT Imaging as a Tool for Studying Mycobacterium tuberculosis Infection and Treatment in Non-human Primates. J Vis Exp doi:10.3791/56375.

93. Moriarty RV, Rodgers MA, Ellis AL, Balgeman AJ, Larson EC, Hopkins F, Chase MR, Maiello P, Fortune SM, Scanga CA, O’Connor SL. 2022. Spontaneous Control of SIV Replication Does Not Prevent T Cell Dysregulation and Bacterial Dissemination in Animals Co-Infected with M. tuberculosis. Microbiol Spectr 10:e0172421.

94. Corbett AJ, Eckle SB, Birkinshaw RW, Liu L, Patel O, Mahony J, Chen Z, Reantragoon R, Meehan B, Cao H, Williamson NA, Strugnell RA, Van Sinderen D, Mak JY, Fairlie DP, Kjer-Nielsen L, Rossjohn J, McCluskey J. 2014. T-cell activation by transitory neo-antigens derived from distinct microbial pathways. Nature 509:361–5.

